# Brain bases for navigating acoustic features

**DOI:** 10.1101/2025.02.10.636597

**Authors:** Alexander J. Billig, William Sedley, Phillip E. Gander, Sukhbinder Kumar, Meher Lad, Maria Chait, Yousef Mohammadi, Joel I. Berger, Timothy D. Griffiths

## Abstract

Whether physical navigation shares neural substrates with mental travel in other behaviourally relevant domains is debated. With respect to sound, pure-tone working memory in humans elicits hippocampal as well as auditory cortical and inferior frontal activity, and rodent work suggests that hippocampal cells that usually track an animal’s physical location can also map to tone frequency when task-relevant. We generated a sound dimension based on the density of random-frequency tones in a stack, resulting in a percept ranging from low- (“beepy”) to high-density (“noisy”). We established that unlike tone frequency, which listeners automatically associate with vertical position, this density dimension elicited no consistent spatial mapping. During functional magnetic resonance imaging, human participants held in mind the density of a series of tone stacks and, after a short maintenance period, adjusted further stacks to match the target (“navigation”). Density of the currently heard sound was represented most strongly in bilateral non-primary auditory cortex, specifically bilateral planum polare, while density of the maintained target was represented in right anterior hippocampus and left inferior temporal gyrus. Encoding and maintenance activity in bilateral hippocampus, inferior frontal gyrus, planum polare and posterior cingulate was positively associated with subsequent navigation success. Bilateral inferior frontal gyrus and hippocampus were among regions with elevated activity during adjustment, compared to a parity-judgment condition with closely matched acoustics and motor demands. Bilateral orbitofrontal cortex was more active when navigation was toward a target density than when participants adjusted density in a control condition with no particular target. We find that self-initiated travel along a non-spatial auditory dimension engages a brain system overlapping with that supporting physical navigation.

**Key Points:**

- Work in rodents suggests that navigation in physical space and the active analysis of sounds share a neural substrate in the hippocampus, supporting the use of common computational mechanisms.
- We examined the human brain system for navigation through an acoustic environment to a remembered target.
- In addition to high-level auditory cortex we demonstrate involvement of the hippocampus in acoustic navigation along with other sites in frontal and cingulate cortex that also support physical navigation.

## Introduction

Finding our way in the physical world is supported by computations involving hippocampus as well as entorhinal, parietal, cingulate, and prefrontal cortices (O’Keefe and Dostrovsky, 1971; Ekstrom et al., 2003; Hafting et al., 2005; Spiers, 2008; Doeller et al., 2010; Bicanski and Burgess, 2020; Basu et al., 2021). Recent theoretical and empirical work suggests that this neural machinery may operate over more general sensory or knowledge spaces (Tavares et al., 2015; Constantinescu et al., 2016; Aronov et al., 2017; Bao et al., 2019; Whittington et al., 2020). In the auditory world a principal organising dimension is frequency. Linguistic and musical communication often depend on generating and making sense of trajectories in frequency, which is mapped tonotopically at multiple levels of the auditory system from the cochlea to sensory cortex. When tone frequency is controlled by a rat to match a target and obtain a reward, neurons tuned to frequency are also found in hippocampus and entorhinal cortex (Aronov et al., 2017). Many of these neurons also represent individual or geometrically related spatial locations (place and grid cells, respectively) during physical navigation, suggestive of a flexible code for traversing different dimensions of experience. Such a correspondence may arise simply because frequency has a natural mapping to spatial location: higher frequencies are implicitly associated with higher points in physical space (Rusconi et al., 2006). It may also be that ensembles of sequentially firing hippocampal cells map progress toward a goal or reward, rather than encoding the particular dimension along which such progress is made (see Zutshi et al., 2025 for an example using tones).

Navigation involves not only adjusting one’s position but also maintaining a representation of the destination. In the spatial domain, hippocampal activity and place coding are more pronounced for navigation toward a target than during aimless movement or foraging (Cornwell et al., 2008; Gulli et al., 2020). Maintaining spatial targets engages frontal sites including ventromedial prefrontal and orbitofrontal cortex (Spiers, 2008; Basu et al., 2021). Holding a tone frequency in mind over seconds engages auditory cortex, frontoparietal regions (particularly inferior frontal gyrus) and hippocampus (Zatorre et al., 1994; Gaab et al., 2003; Kumar et al., 2016; Uluç et al., 2018; Kumar et al., 2021; Deutsch et al., 2023). A similar network supports maintenance of more complex auditory objects, such as spectral ripples or tones defined by a combination of frequency and either location or presentation rate (Czoschke et al., 2021; Ahveninen et al., 2023; Pomper et al., 2023; Uluç et al., 2023). We designed a stimulus and task that allowed us to probe the neural substrates of what we define as non-spatial “navigation” toward a spectrotemporally complex target sound in humans.

We used successions of tone stacks that varied in the number of concurrently presented tones of random frequency (Figure 1A). This gave rise to a percept varying from sparse (“beepy”) to dense (“noisy”). Using objective and subjective behavioural measures we established that this acoustic dimension does not consistently elicit a spatial association. We then had participants undergo functional magnetic resonance imaging (fMRI) while performing tasks that separately tapped processes of working memory and navigation in this auditory dimension. The main condition of interest involved holding in mind a series of tone stacks of fixed density, then adjusting a new set of stacks to match the target. In order to establish whether any coding for density was distinct from proximity to the target we used different starting and target densities in different trials. In control conditions we removed one or both of (a) the working memory component by having participants randomly adjust tone stacks with no particular target, and (b) the navigation component by having participants respond as to whether each in a series of spoken numbers was odd or even while listening passively to the tone stacks. As far as possible we matched the acoustics and number of button presses across conditions.

**Figure 1:**
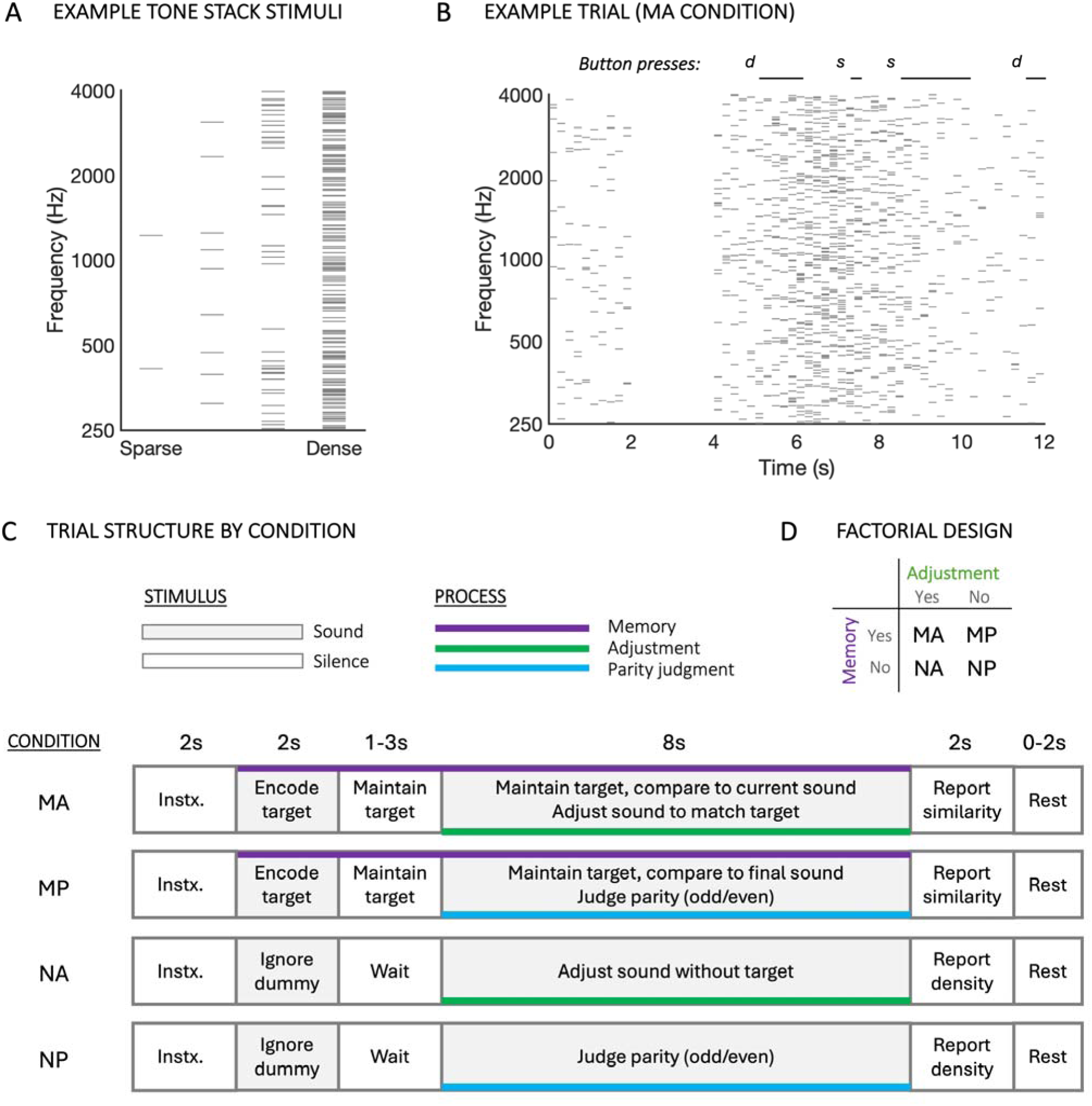
Stimuli and experimental design. **A)** Four example tone stacks ranging in spectral density from sparse (“beepy”, left) to dense (“noisy”, right) based on the number of pure tone components. In the pilot behavioural experiment participants heard two such stacks and judged whether the second was beepier or noisier than the first. **B)** Example of an MA (memory adjust) trial in the fMRI experiment, showing target sound (starting at 0s), maintenance period (starting at 2s), and adjustment period (starting at 4s). Tone components are shown as short horizontal grey lines. Button presses (reflecting attempts to match the target density) are shown as ‘d’ (make denser) and ‘s’ (make sparser) with black horizontal lines representing their durations. **C)** Structure of trials in each of the four conditions. Conditions with labels containing ‘M’ involved maintaining a target density in <u>m</u>emory (purple lines) and reporting its similarity to that at the end of a subsequent series of stacks; conditions with labels containing ‘N’ did <u>n</u>ot involve target maintenance - instead participants reported how beepy or noisy the sound at the end of the series of stacks was. Conditions with labels containing ‘A’ (green lines) involved <u>a</u>djusting the ongoing series of stacks (either to match the target density in the MA condition, or at will in the NA condition); conditions with labels containing ‘P’ (blue lines) instead required participants to report the <u>p</u>arity of each a series of spoken digits (i.e. whether it was odd or even). Tone stacks and digits were present in all conditions, including when task-irrelevant. **D**) Factorization of the design into four conditions based on the presence or absence of memory and adjustment processes. Instx. = Instructions, MA = Memory Adjust, MP = Memory Parity, NA = No-memory Adjust, NP = No-memory Parity.

Regardless of task, we expected the spectral density of sounds that were matched for overall spectrum to be represented in higher-order auditory cortex on the superior temporal plane. Based on previous imaging work with a range of acoustic stimuli we hypothesized that auditory cortex, inferior frontal gyrus and hippocampus would be active when such sounds were maintained as targets. If mechanisms of spatial navigation and auditory analysis overlap in humans as they do in rodents we anticipated hippocampal activation and representation of the auditory space during adjustment, but not for the control task. We also expected any such hippocampal engagement to be more pronounced when auditory adjustment was toward a stored target than when it was aimless.

## Materials and Methods

### Stimuli

The primary experimental stimuli were 190-ms tone stacks (with 2-ms cosine on- and offset ramps) interspersed with 10 ms of silence. Varying the number of concurrent pure tones in a stack gave rise to a “density” percept described to participants as ranging from “beepy” to “noisy” (see Figure 1A). In the main experiment each stack contained between 2 and 196 tones, this value taking one of 26 logarithmically spaced levels. In the pilot study, an upper density limit of 400 tones was used, and density levels were determined by the staircase described in “Behavioural pilot experimental procedure”.

In both the pilot and the main experiment, tone frequencies were randomly drawn from a logarithmic distribution between 250 and 4000 Hz. Tone stacks were equated for root-mean-square intensity, which gave a reliable loudness match during piloting. We also created recordings of spoken digits from 1 to 10 using a text-to-speech tool (notevibes.com; voices: Daniel, Gabriela; speed: 1.6). These were trimmed to have duration below 400 ms and normalised for root mean square intensity.

### Participants

Ten individuals (aged 21-40 years, median 26.5 years, 4 males, 6 females) with self-reported normal hearing took part in the behavioural pilot experiment. A different group of 30 right-handed individuals with no reported neurological or hearing impairment took part in the fMRI experiment. Data from 25 of these (aged 19-40 years, median 24 years, 12 males, 13 females) were included in the analysis. Of the remaining five participants, one did not complete the experiment, two performed more than two standard deviations below the group mean in the adjustment task (see fMRI experiment procedure), and three (including one of the poor performers) moved more than 3 mm (one voxel) in the scanner. Participants were not selected on the basis of musical training in either experiment. The study was approved by the University College London Research Ethics Committee. Informed consent was obtained from all participants, who were compensated for their time and travel.

### Behavioural pilot experimental procedure

The purposes of the behavioural pilot experiment were to establish (1) how well participants could discriminate different levels of tone stack density and (2) whether reaction times were affected by the layout of response keys, potentially reflecting an implicit mapping between density and physical space. After familiarization with the density feature and instruction that avoided language relating to space, participants heard pairs of 200-ms tone stacks separated by 1s of silence. Participants had to respond whether the second stack was “beepier” or “noisier” than the first, as quickly and accurately as possible. Different pairs of keys were mapped to “beepier” and “noisier” responses in eight blocks. In two blocks the “down” arrow was used for a “beepier” response and the “up” arrow for a “noisier” response; in another two blocks this was reversed; in two further blocks the “left” arrow key was assigned to “beepier” and the “right” arrow to “noisier”; in a final two blocks this was reversed. One of the two stacks in each trial (selected at random) took one of four logarithmically spaced reference density levels (3.5, 9.1, 25.2, 66.5 tone components/octave). The density difference between the two stacks in a trial was adaptively tracked using a two-down, one-up staircase procedure converging on the participant’s 70.7% correct threshold (averaging over the final two of six reversals, step size 1/3 until second reversal then 1/6). Two adaptive tracks, each featuring a particular reference density, were run in parallel in each of the eight blocks, resulting in sixteen tracks. The assignment of reference density levels to button mappings was counterbalanced over participants, as was the block order. Accuracy feedback was provided after each response.

### Online training for fMRI participants

To ensure that participants were proficient at the main experimental tasks before scanning, we provided online training (presented in Gorilla with custom JavaScript code) during the preceding week. Participants were familiarized with the density manipulation and then practised each element of each task (see next section) with feedback, before performing several trials of the full experiment. Further training trials were administered in person immediately prior to scanning.

### fMRI experimental procedure

Participants took part in four experimental runs in the scanner, each lasting 8.5 minutes and comprising 28 trials. Each run contained seven trials of each of four conditions, which separately manipulated the presence of memory and adjustment task components while balancing acoustics and button presses. Figure 1C illustrates the trial structure for each condition, and videos of example trials are available at in Supporting Information. In both the “Memory Adjust” (MA) and “Memory Parity” (MP) conditions, participants held in mind the density of a target sound over a maintenance period. In the MA condition participants then adjusted the density of a series of stacks (with starting density chosen randomly) to match the target density (Figure 1B). At the end of the trial, they rated how close in density the final sound was to the target. In the MP condition after hearing the target they performed an “odd/even” task on a series of spoken digits. They then turned their attention to the end of an ongoing series of tone stacks and rated its similarity to the initial target. The “No-Memory Adjust” (NA) and “No-Memory Parity” (NP) conditions did not include a target. In the NA condition, after hearing a series of stacks of mixed density, participants adjusted a new series of stacks at will. They then rated how “beepy” or “noisy” the final adjusted stack was. The NP condition also began with a series of stacks of mixed density, was followed by the “odd/even” task on spoken digits and ended with a rating of how “beepy” or “noisy” the final stack of an ongoing series was. Trials began with two seconds of silence while instructions were displayed on screen. These varied with condition, included a reminder of the available response options, were shown in black text on a light grey background, and covered all three phases of the trial (see Table 1). Once the sounds started, the instructions for the relevant phase were highlighted in white.

**Table 1.** Participant instructions for each condition.

| Condition code | Condition description | Phase 1 instructions | Phase 2 instructions | Phase 3 instructions |
| --- | --- | --- | --- | --- |
| MA | Memory, Adjust | Store | Adjust | Compare |
| MP | Memory, Parity | Store | Odd/Even | Compare |
| NA | Non-memory, Adjust | Ignore | Adjust | Beepy/Noisy |
| NP | Non-memory, Parity | Ignore | Odd/Even | Beepy/Noisy |

The initial sound in all conditions was a series of ten 190-ms stacks, each followed by 10 ms of silence. In the MA and MP conditions the density of these stacks was set to a fixed target value, the specific frequencies varying from stack to stack over the resulting 2-s target. In the NA and NP conditions these initial stacks instead alternated between the lowest- and highest-density values. This was to emphasise that they formed a “dummy” sound that was to be ignored, while matching the average spectrum and density across conditions. In all conditions, the 2-s sound was followed by 1-3 s of silence, sampled from a rectangular distribution – in the MA and MP conditions this silence acted as a working memory maintenance phase. Over the next 8 s two simultaneous sound streams were heard: (1) a series of 40 stacks and (2) between one and six spoken digits. The value of each digit was randomly selected between 1 and 10 and spoken by either a male or female voice, also randomly selected. Digits could appear in slots beginning at one-second intervals from second two of the eight-second phase, leaving the first and final second free of digits such that only the tone stacks were audible. In the MA and NA (adjustment) conditions participants ignored the digits and directed attention to the tone stacks throughout. They held down one right-hand button to increase the density of the subsequent tone stacks, and another right-hand button to reduce it - a new random set of frequency components was selected for each tone stack regardless of whether the density changed. In the MA condition they were instructed to try to match the target as closely as possible, using as much of the eight seconds as needed. In the NA condition they adjusted the sounds at will and were asked to use a similar number of presses as in the MA trials.

In the MP and NP (parity) conditions, participants attended to spoken digits and responded via one of two right-hand buttons as to whether each digit was odd or even (a similar task has served as an appropriate baseline for hippocampal studies in the visual domain; Stark and Squire, 2001). To try to match the number of button presses across conditions we set the number of digits presented in each trial to that during the previous MA or NA trial plus the mean number of digits for which no response was made across all previous MP and NP trials (or 0.5 when no such trials were available at the start of a block), rounded up or down (at random) to the nearest integer. For the last second of this 8-s section of MP and NP trials, participants redirected their attention to the final few tone stacks. To match acoustics across conditions, in these MP and NP trials the densities of the ongoing tone stacks matched those generated in the most recent MA or NA trial respectively. Identity, speaker, and timing of (ignored) digits in the MA and NA trials were identical to those in the most recent MP or NP trials respectively (or, when no previous trials were available at the start of a block, to those in a randomly generated dummy trial). In the MA and MP conditions, the target density and starting density of the 8-s tone series of stacks for each trial were randomly drawn from a predetermined set of 28 pairs that would generate a range of expected trajectories spanning the density space with equal sampling of high- and low-density sounds. In the NA and NP conditions the starting density of the 8-s series of tone stacks for each trial was drawn from the same set of 28 pairs (with the target density disregarded).

The final phase of each trial required a judgment of either how different the density of the final tone stack was to that of the target (MA and MP trials; response options: “Very similar”, “Somewhat similar”, “Somewhat different”, “Very different”) or how beepy/noisy the final tone stack was (NA and NP trials; response options: “Very beepy”, “Somewhat beepy”, “Somewhat noisy”, “Very noisy”). Participants had three seconds to respond using one of four left-hand buttons. After a pause of between 0 and 2 s (sampled from a rectangular distribution) the next trial began. No feedback was provided. The 28 trials in a block were grouped into seven sets of four (not made explicit to participants), each set containing one trial of each condition. Ordering within each set was pseudo-randomized, with the constraint that the MA trial appeared earlier than the NA trial, and the MP trial appeared earlier than the NP trial. This ensured that the 8-s series of tone stacks in the NA and NP trials could be matched to those in the previous MA or NA trials, while preventing two trials of the same condition occurring in succession.

The first two experimental blocks were separated by a break of up to two minutes. These were followed by field mapping and anatomical scans (apart from for one participant whose anatomical scan was obtained at the start of the session), lasting a total of around ten minutes. The final two experimental blocks followed and were also separated by a break of up to two minutes. Sounds were presented through an Edirol UA-4FX (Roland Corporation, Shuzuoka) USB soundcard and Ear-Tone Etymotics tube earphones (Etymotic Research, Inc.) with foam tips. Levels were tested and adjusted at the start of the session to ensure that both the tone stacks and spoken digits were audible over the scanner noise. Instructions were displayed via an Epson EB-L1100U projector and viewed with a head-coil-mounted mirror. The experiment was controlled using Psychtoolbox and custom MATLAB code.

### MRI data acquisition

Data were collected using a 3-Tesla Siemens MAGNETOM Prisma MR scanner (Siemens Healthcare) with a 32-channel receive head coil at the Wellcome Centre for Human Neuroimaging (London, UK). Whole-brain T2*-weighted functional images were acquired using 3D echo-planar imaging with field of view 192 x 192 x 144 mm, voxel size 3 mm isotropic, 48 transverse slices, 3D acceleration factor 2, repetition time (TR) 44 ms, volume acquisition time 1056 ms, anterior-to-posterior phase encoding, and bandwidth 2298 Hz/Px. The accelerated functional images were reconstructed using a motion-robust higher-order sensitivity encoding approach (https://github.com/fil-physics/gadgetron-matlab/tree/master/FIL-recon). Respiratory signals were obtained with a breathing belt. Field maps (double-echo gradient echo sequence with field of view 192 x 192 x 144 mm, voxel size 3 x 3 x 2 mm, 64 transverse slices, TR 1020 ms, short echo time (TE) 10.00 ms, long TE 12.46 ms, flip angle 90°) and a whole-brain T1-weighted anatomical image (MPRAGE, field of view 256 x 256 x 176 mm, voxel size 1 mm isotropic, 176 transverse slices, GRAPPA acceleration factor 2, TR = 2530 ms, TE = 3.34 ms) were acquired during the same scanning session.

### MRI preprocessing

MRI data were processed in SPM12 (Wellcome Centre for Human Neuroimaging). Each participant’s functional images were unwarped using their field map and realigned to the first volume. Functional and anatomical images were coregistered to the mean functional image. For the univariate analyses, the coregistered images were normalized to the SPM12 template (avg305T1) in MNI space and the functional images were spatially smoothed using a Gaussian kernel with 6 mm full-width at half-maximum.

### Behavioural data analysis

For the behavioural experiment, participants’ just-noticeable-differences (JNDs) were obtained as a function of density. Reaction times were log-transformed and analysed as a function of button mapping using a repeated-measures 4-way ANOVA. For the fMRI experiment, navigation success was scored using the absolute number of steps in density space between the final adjusted sound and the target. Parity judgments were scored by dividing the number of matches by the larger of two values: the number of digits presented or the number of responses made. When determining matches, the series of responses in a trial was allowed to be dislocated by an extra or missing response. Accuracy of responses to the final question in each trial was assessed by calculating the Spearman’s rank correlation of the responses (1-4) with the true values (closeness in density for MA and MP conditions; actual density for NA and NP conditions), after replacing missing responses with a value of 2.5 (the midpoint of both rating scales). Proportions were compared following arcsine square root transformation, and correlation coefficients following the Fisher *z*-transformation.

### Univariate fMRI analyses

In a first level analysis a General Linear Model (GLM) was created for each participant’s fMRI time series data. Regressors of no interest were included for motion and respiration. Task phases of interest were modelled as boxcar functions and convolved with the canonical haemodynamic response function. A high pass filter of 1/230 Hz (corresponding to twice the longest interval between trials of the same condition) was applied to remove low frequency noise, and SPM12’s “FAST” option (appropriate for rapid imaging sequences; Corbin et al., 2018) was used to model temporal correlations. To distinguish activity relating to the different task components we first modelled the whole trial period and tested for main effects of memory [contrast: (MA+MP)-(NA+NP)] and adjustment [contrast: (MA+NA)-(MP+NP)], and for their interaction [contrast: (MA-MP)-(NA-NP)]. In a control analysis we modelled these effects only during the portion of the MA trials during which adjustment responses were made (and matching portions of trials in other conditions). Effects of density (all trials), distance from target (MA trials), time in trial (MA trials), and performance in adjustment (MA trials) and parity judgment (MP trials) tasks were examined by entering centered and orthogonalized linear parametric modulators into the GLM over the relevant time intervals.

For each comparison of interest, participants’ contrast images were entered into a second level analysis, with one-sample *t*-tests conducted at the group level. Statistical parametric maps were obtained and thresholded using a family-wise error- (FWE-) corrected *p*-value of .05, and a minimum cluster size matching the “expected voxels per cluster” value in SPM for the corresponding height threshold. Additional tests with small volume corrections were conducted in three a priori regions of interest, namely bilateral auditory cortex (voxels assigned at least 20% probability of lying in Heschl’s gyrus, planum polare, planum temporale, or superior temporal gyrus in the Harvard-Oxford Cortical Structural Atlas), bilateral inferior frontal gyrus pars opercularis and pars triangularis (voxels assigned at least 20% probability of lying in one of these two regions in the Harvard-Oxford Cortical Structural Atlas), and bilateral hippocampus (based on segmentation of the average T1w image across participants using the Automatic Segmentation of Hippocampal Subfields (ASHS) software package, Yushkevich et al., 2014).

### Multivariate fMRI analyses

Multivariate analyses were performed across participants, an approach that tests for not only the amount of information encoded in a given region but also the consistency of such a voxel-level information code. It yields results that are more generalisable than within-participant analyses and can detect smaller effects when they are consistent (Wang et al., 2020). To identify multivoxel representations of density and adjustment direction, the unsmoothed images (normalised to the *avg305T1* SPM12 template in MNI space for the across-participant analysis only) were entered into further GLMs. One contained regressors for lower-versus higher-density sounds (splitting the 26 density levels into two groups) and used trials of all conditions. Another contained regressors for navigation in the direction of decreasing versus increasing density and used MA and NA trials. In both models these binary labels were applied to each stack during the adjustment period and could therefore change within a trial. A further analysis used the same approach to find distances between representations of lower-versus higher-density targets across the whole trial (encoding, maintenance and adjustment/parity judgment periods) in MA and MP conditions. In all GLMs the same regressors of no interest were entered as described in Univariate fMRI analyses above. Multivariate analysis was performed using The Decoding Toolbox (Hebart et al., 2015) to obtain cross-validated whitened Euclidean (“crossnobis”) distances between neural representations corresponding to the two category labels in a given model. Neural representations were beta weights scaled separately for each subject using residuals from the corresponding GLM with the “Ledoit-Wolf retain variances” option selected. A spherical volumetric searchlight with 4 mm radius was used over the whole brain. Leave-one-participant-out cross-validation was applied and the lower diagonal values of the dissimilarity matrices extracted for each cross-validation fold. Each cross-validation fold’s distance image corresponded to testing a single participant on a model trained on all the others. The corresponding 25 images were spatially smoothed using a Gaussian kernel with 6 mm full-width at half-maximum. These were entered into second-level analyses, which proceeded as described in Univariate fMRI analyses.

## Results

### Tone stack density: discrimination and associations with physical space

In the initial behavioural experiment, just-noticeable differences for density increased with density [*t*(9) = 13.717, *p* < .001, *d* = 4.338; Figure 2A]. For this reason, density values in the fMRI experiment were logarithmically spaced to ensure steps along the density scale were similarly discriminable. Reaction times when comparing the density of two tone stacks was not affected by the spatial mapping of response keys [*F*(3, 27) = 0.549, *p* = .653, *η*^2^_p_ = .057; Figure 2B]. When prompted, twelve of the participants in the fMRI experiment reported forming a spatial association with the stimuli (nine associated beepier sounds with “left”, one with “right”, and two with “up”). To confirm that the group neural results were not driven by these spatial associations we also report the relevant analyses by excluding the corresponding participants, obtaining qualitatively similar results.

**Figure 2:**
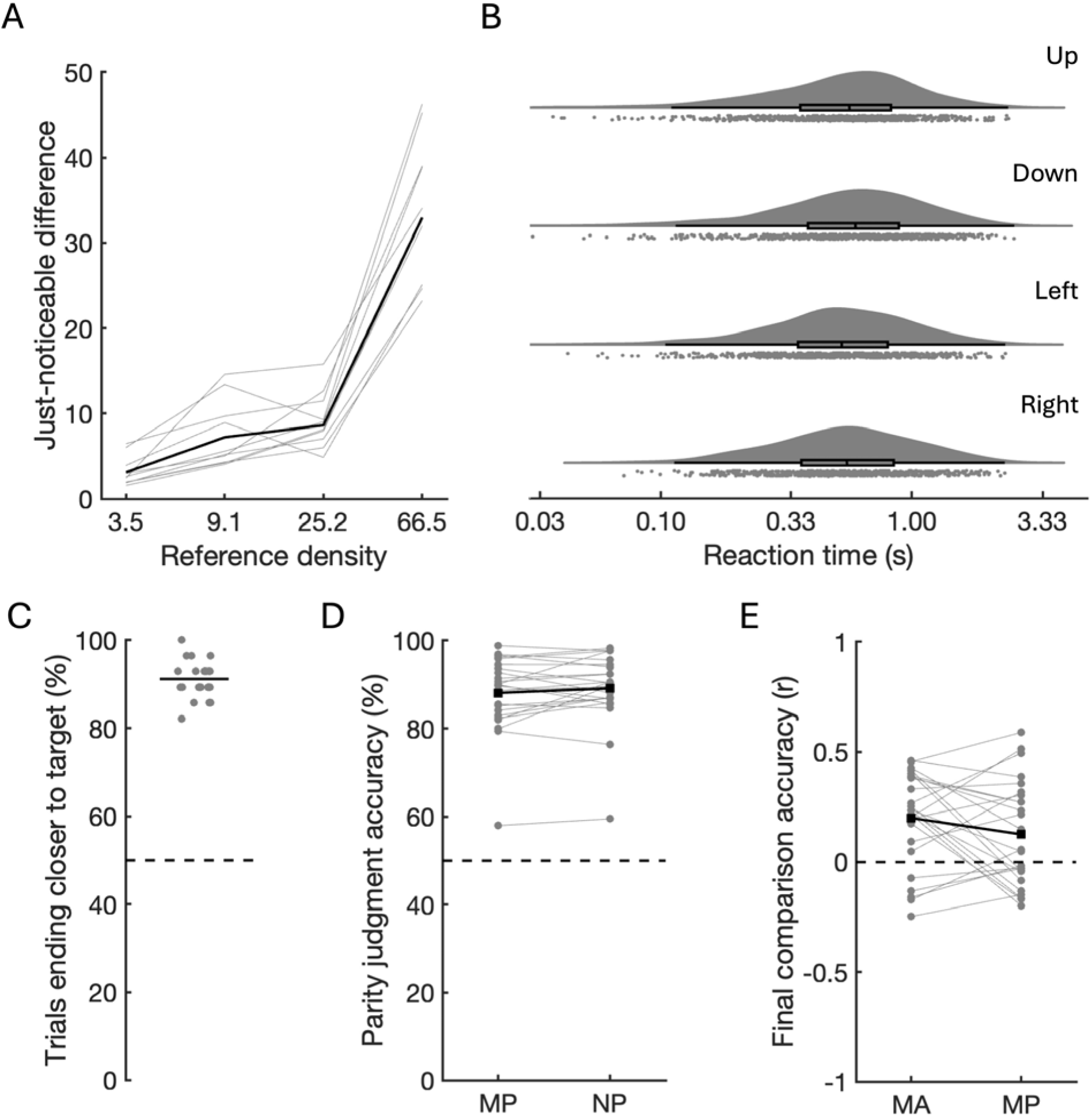
Behavioural results. **A)** Just-noticeable differences in density increase as a function of reference density; values are the number of tone components per octave. Grey lines: individual participants in the behavioural pilot experiment. Black line: group mean. **B)** Raincloud and box-and-whisker plots of reaction times for correct trials in the behavioural pilot experiment. Each plot corresponds to a different button mapping (the arrow key that was mapped to “beepier” responses is shown; the opposite arrow key was mapped to “noisier” responses). Distributions did not differ by condition. **C)** Percentage of MA trials in the main experiment in which participants (grey dots) adjusted the ongoing density to be closer to the target than before adjustment. Black line represents mean performance, and dotted line represents 50% chance performance. **D)** Percentage accuracy of parity judgments, which did not differ significantly between MP (left) and NP (right) conditions. Joined grey dots represent individual participants. Joined black squares represent group mean performance and dotted line represents 50% chance performance. **E)** Correlation between actual and reported target-versus-final density similarity, which did not differ significantly between MA (left) and MP (right) conditions. Joined grey dots represent individual participants. Joined black squares represent group mean performance and dotted line represents chance performance.

### Behavioural performance in the scanner

In the MA (memory adjustment) condition of the main experiment, participants finished closer to the target than they started in 94% of trials on average (range: 82-100%; Figure 2C), with a mean final distance of 16% of the density scale (range: 9-33%). Participants tended to undershoot the target, that is to remain on the same side of the target on the density scale as they began; on average this occurred in 70% of trials (range: 32-86%; Supplemental Figure S1). Performance did not differ between participants who reported forming spatial associations and those who did not [*t*(23) = 0.503, *p* = .620, *d* = .100]. Parity judgment performance did not differ between the memory and non-memory conditions [MP and NP; *t*(24) = 1.132, *p* = .269, *d* = 0.227; Figure 2D], with an average score of 89% (range: 59-98%). Accuracy in comparing the target density to the final density did not differ significantly between the MA and MP conditions [*t*(24) = 1.475, *p* = .139, *d* = 0.298; Figure 2E], while accuracy in reporting the final density was significantly greater in the NA than the NP condition [*t*(24) = 2.441, *p* = .022, *d* = 0.584].

### Matching of motor activity across conditions

There was a significant effect of condition on the number of presses despite our attempt to eliminate this potential confound [*F*(3, 72) = 10.256, *p* < .001, *η*^2^_p_ = .299]. Although the majority of participants were well matched (on average a difference of less than one press per trial between pairs of conditions of interest, namely MA versus NA, and MP versus NP), eight participants had more than one extra press on average in adjustment trials (MA, NA) compared to parity trials (MP, NP), and six participants had more than one extra press on average in non-memory trials (NA, NP) compared to memory trials (MA, MP). Eleven participants met both criteria. To confirm that the group neural results were not confounded by activity relating to motor preparation or execution we also report the relevant analyses excluding the corresponding participants, obtaining qualitatively similar results.

### Neural responses to sound and representation of tone stack density

Sites that were more active during sound compared to silence were bilateral auditory cortices in Heschl’s gyrus, planum temporale, and posterior superior temporal gyrus, along with parietal, somatomotor, frontal, parietal and cerebellar areas (see Table 2). This contrast did not reflect purely sensory processes since cognitive and motor demands also differed when sound was presented compared to when it was not. During the adjustment phase (or equivalent period in the parity judgment task), activity in a more restricted set of brain regions, shown in Figure 3A and Supplemental Table S1, was parametrically modulated by the density of the tone stacks. Activity was greater for the sparser (less noise-like) tone stacks - throughout much of bilateral auditory cortex. During the same phase, multivoxel patterns for sparser versus denser sounds were also most distinct at a subset of these sites, most markedly in bilateral planum polare (Figure 3B, Table 3). After small volume correction, a significant representation of the currently presented density was also found in right inferior frontal gyrus (pars opercularis; Supplemental Table S2), but not hippocampus.

**Figure 3:**
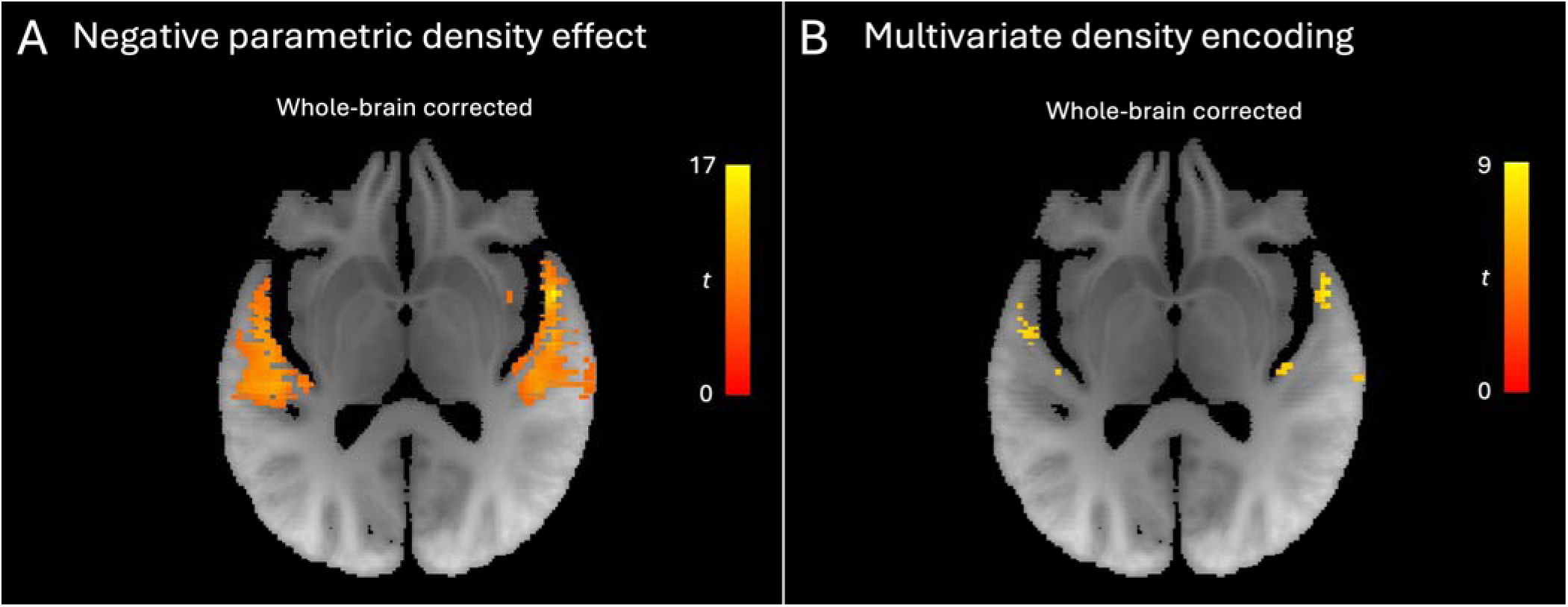
Representation of tone stack density. **A)** Tilted axial slice (aligned with the superior temporal plane) showing negative parametric effect of density in auditory cortex, i.e. a lower univariate response to progressively denser tone stacks. **B)** Tilted axial slice (aligned with the superior temporal plane) showing multivariate distance (transformed to *t*-score) between representations of sparser and denser tone stacks in auditory cortex. All clusters significant at *p* < .05 (familywise error-corrected, whole brain).

**Table 2.** Clusters more active during sound presentation than silence.

| Regions | Voxels | Peak MNI coordinates |  |  | Peak t-value |
| --- | --- | --- | --- | --- | --- |
| Right planum temporale, Heschl's gyrus, superior temporal gyrus (posterior) | 407 | 56 | -14 | 6 | 10.29 |
| Left planum temporale, Heschl's gyrus, superior temporal gyrus (posterior) | 333 | -54 | -28 | 10 | 9.65 |
| Left supplementary motor area, superior frontal gyrus | 131 | -6 | 8 | 54 | 9.45 |
| Left precentral gyrus, postcentral gyrus | 174 | -40 | 18 | 60 | 8.89 |
| Right superior frontal gyrus, supplementary motor area | 108 | 6 | 10 | 58 | 8.11 |
| Left supramarginal gyrus (anterior) | 72 | -52 | -32 | 50 | 7.79 |
| Left precentral gyrus | 16 | -56 | 6 | 18 | 7.55 |
| Right inferior frontal gyrus (pars opercularis) | 14 | 52 | 10 | 10 | 6.85 |
| Left precentral gyrus | 7 | -28 | -6 | 62 | 6.49 |
| Right cerebellum (lobe V) | 5 | 20 | -52 | -24 | 6.19 |

**Table 3.** Clusters with multivoxel representation of density.

| Regions | Voxels | Peak MNI coordinates |  |  | Peak t-value |
| --- | --- | --- | --- | --- | --- |
| Right planum polare | 68 | 56 | 6 | 0 | 8.89 |
| Left planum polare | 74 | -44 | -20 | -4 | 8.00 |
| Right planum polare | 46 | 44 | -16 | -2 | 7.42 |
| Right Heschl's gyrus | 20 | 42 | -22 | 10 | 7.33 |
| Right superior temporal gyrus (posterior) | 8 | 46 | -22 | -2 | 7.06 |
| Left Heschl's gyrus | 3 | -42 | -22 | 8 | 6.97 |
| Right superior temporal gyrus (anterior) | 14 | 64 | -2 | -8 | 6.91 |
| Left insular cortex | 11 | -32 | -28 | 14 | 6.88 |
| Left central opercular cortex | 2 | -50 | -6 | 6 | 6.86 |
| Left superior temporal gyrus (posterior) | 2 | -68 | -24 | 2 | 6.85 |
| Right superior temporal gyrus (posterior) | 4 | 68 | -24 | 6 | 6.83 |
| Left superior temporal gyrus (anterior) | 6 | -60 | -12 | -2 | 6.70 |
| Right inferior colliculus | 2 | 12 | -28 | -10 | 6.65 |
| Left insular cortex | 2 | -38 | 2 | -10 | 6.66 |

### Memory-related neural activity

Next we contrasted activity in the two conditions in which participants held a target density in mind (MA and MP) versus those with no such target (NA and NP). We considered activity throughout the trial, from the start of the instructions to the end of the final report (a period lasting an average of 16 s and corresponding to all but the final column in Fig. 1C). This revealed memory-related activity in bilateral anterior insula, as well as frontal pole, middle frontal gyrus, and cerebellum (Figure 4 middle, right; Table 4). Activity levels for each condition averaged across voxels in each significant cluster are shown in Supplemental Figure 2. After small volume correction, memory-related activity was also found in bilateral inferior frontal gyrus (pars opercularis and pars triangularis, Figure 4 (left), Supplemental Table S3), but not hippocampus. In addition to the density representations of sounds as they were presented (described in the previous section), we found the target density represented throughout MA and MP trials in left inferior temporal gyrus (after whole-brain correction) and right anterior hippocampus (after small volume correction), as shown in Fig. 5 and Supplemental Tables S4 and S5.

**Figure 4:**
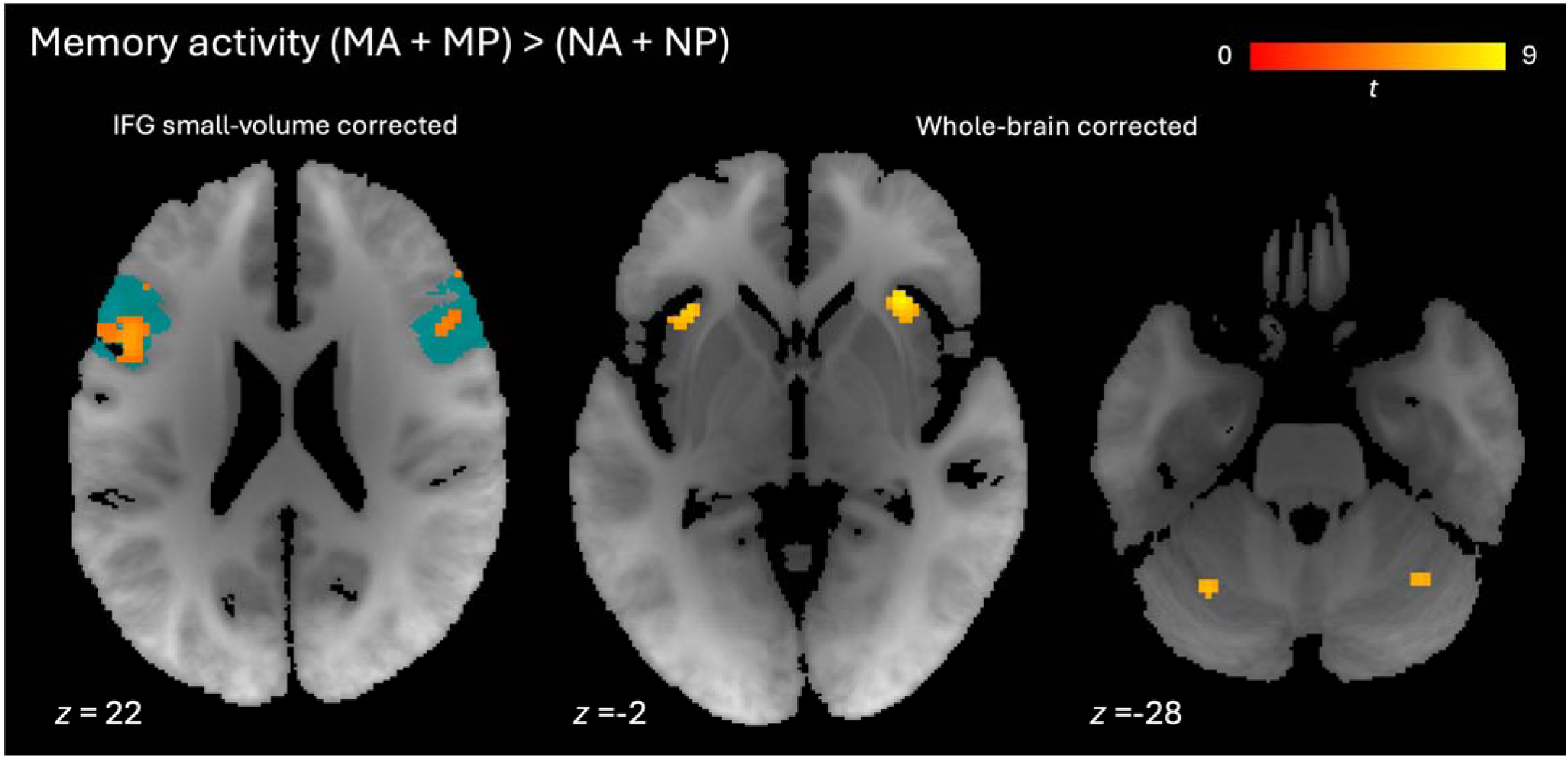
Activity associated with target memory. Axial slices showing clusters with greater activity for MA and MP conditions than NA and NP conditions. **Left**: Inferior frontal gyrus clusters significant at *p* < .05 (familywise error-corrected, small volume for bilateral inferior frontal gyrus). **Middle**: Anterior insular clusters significant at *p* < .05 (familywise error-corrected, whole brain). **Right**: Cerebellar clusters significant at *p* < .05 (familywise error-corrected, whole brain). Red-yellow gradient represents *t*-score. Blue indicates regions of interest for small volume corrections. IFG = Inferior frontal gyrus.

**Figure 5:**
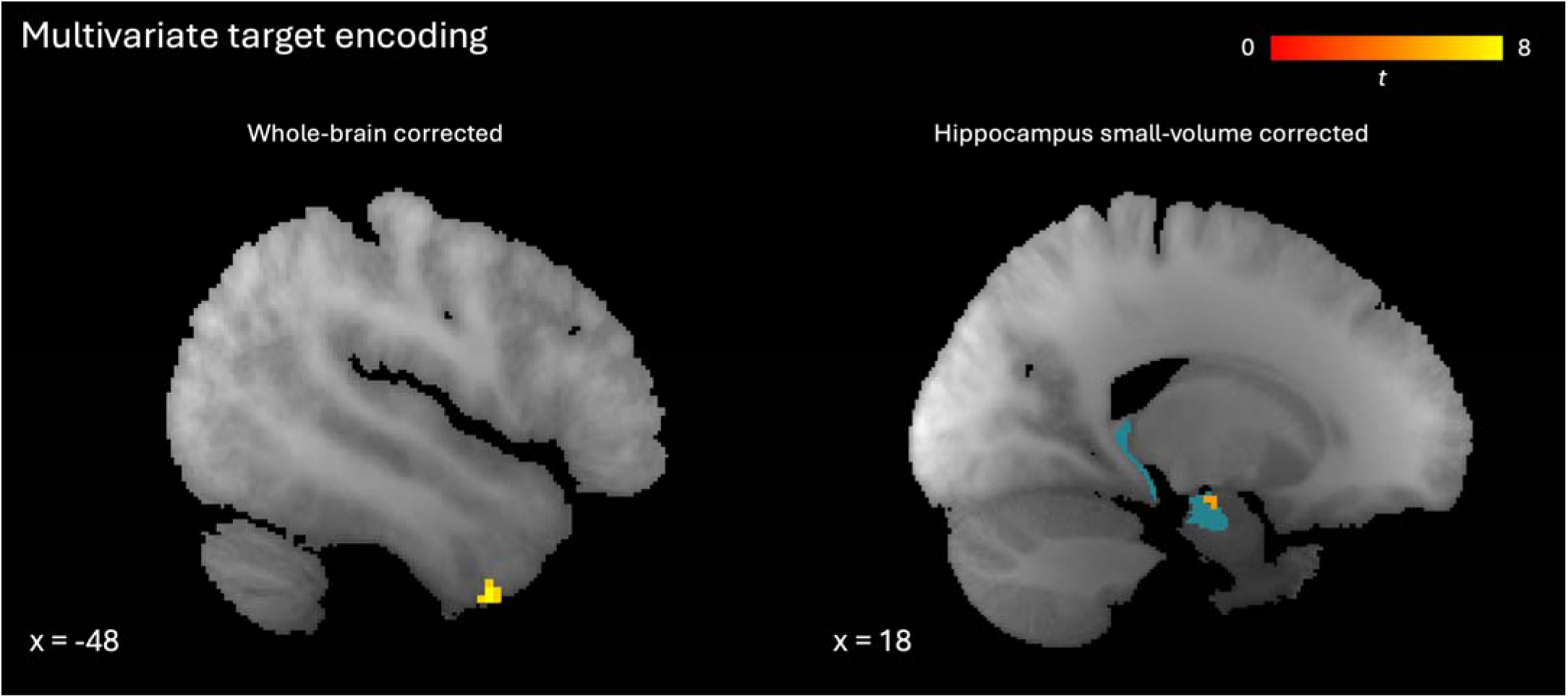
Representation of target density. Sagittal slices showing multivariate distance (transformed to *t*-score) between representations of sparser and denser target tone stacks (MA and MP conditions). **Left**: Left inferior temporal gyrus cluster significant at *p* < .05 (familywise error-corrected, whole brain). **Right**: Right anterior hippocampal cluster significant at *p* < .05 (familywise error-corrected, small volume for bilateral hippocampus). Red-yellow gradient represents *t*-score. Blue indicates region of interest for small volume correction.

**Table 4.** Clusters more active during memory than non-memory conditions.

| Regions | Voxels | Peak MNI coordinates | | | | Peak $t$-value |
| --- | --- | --- | --- | --- | --- | --- |
| Right insular cortex | 58 | 30 | 24 | 0 |  | 8.92 |
| Right frontal pole | 30 | 46 | 40 | 28 |  | 8.23 |
| Left insular cortex | 43 | -32 | 20 | -4 |  | 7.50 |
| Left middle frontal gyrus | 6 | -44 | 2 | 54 |  | 6.90 |
| Left cerebellum (lobe VI) | 13 | -30 | -64 | -28 |  | 6.76 |
| Right cerebellum (crus I) | 11 | 34 | -62 | -28 |  | 6.46 |

Clusters significant at $p < .05$ (FWE-corrected), minimum cluster size: 5 voxels

### Adjustment-related neural activity

The contrast of adjustment conditions (MA + NA) minus non-adjustment conditions (MP + NP) revealed activity in our regions of interest of hippocampus and inferior frontal gyrus after small volume correction (Figure 6; Supplemental Table S6). The corresponding clusters remained significant when excluding participants who reported noticing a spatial mapping of density (Supplemental Table S7) or for whom button presses were not well matched across conditions (Supplemental Table S8). At the whole brain level, additional frontal, parietal, insular, cingulate and cerebellar sites also showed adjustment-related activity (Table 5, Figure 7). The more anterior of the adjustment-related hippocampal clusters overlapped with that at which the target density was represented (compare Figure 5 right and Figure 6 left). However, whether these functions (target storage and navigation) are subserved by the same circuits in anterior hippocampus is unclear due to the following additional analysis. We considered the fact that even when button presses were comparable across conditions, participants could choose to stop adjusting before the end of an adjustment trial whereas they needed to remain alert throughout parity judgment trials, even if they made no further presses. It is therefore possible that some differences were driven by task-negative (e.g. mind-wandering) activity toward the end of adjustment trials. In an analysis including only the interval in a given MA or NA trial during which adjustments actually occurred and using the same intervals in matching MP and NP trials, the more anterior of the hippocampal clusters in Figure 5 was no longer significant for the adjustment contrast (Supplemental Table S9).

**Figure 6:**
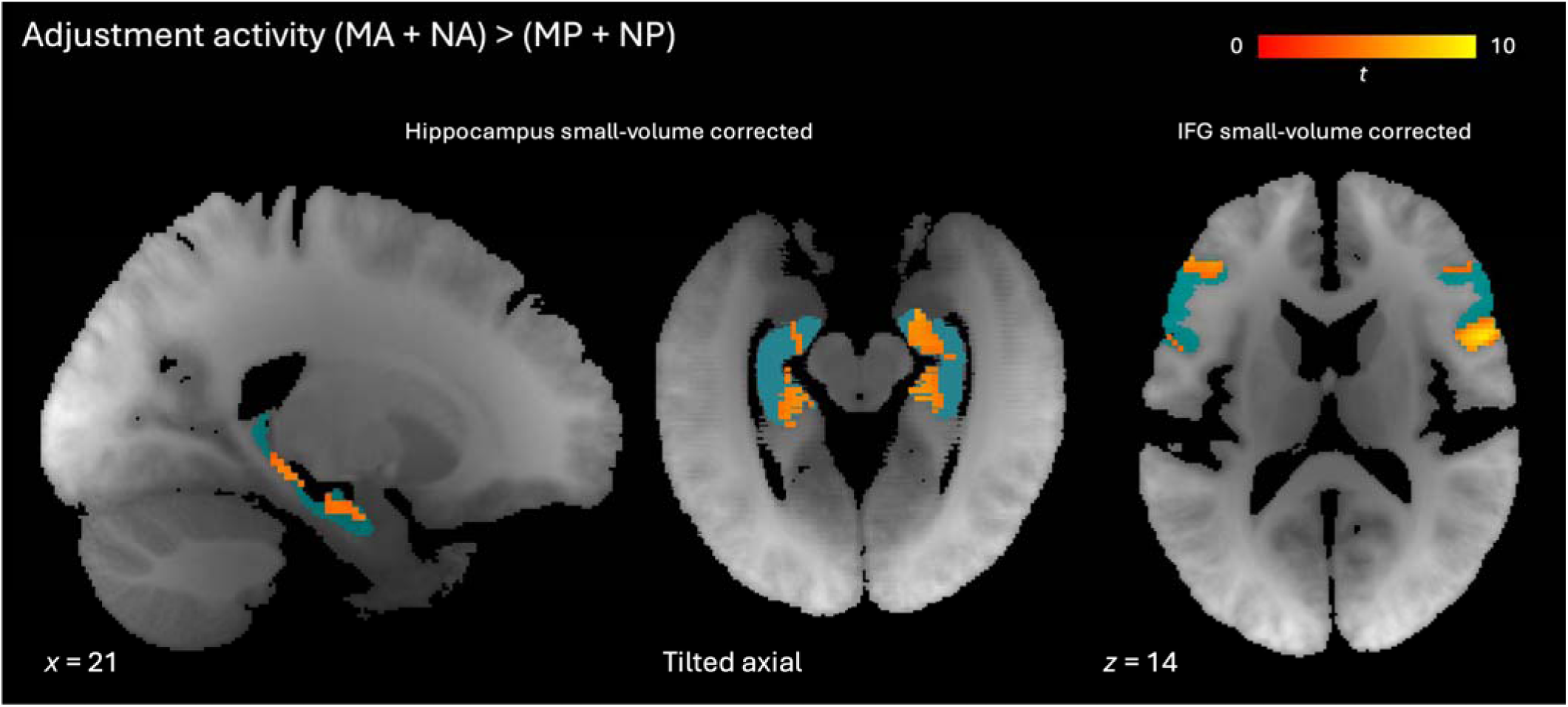
Activity associated with density adjustment in regions of interest. Clusters showing greater activity for MA and NA conditions than MP and NP conditions. **Left**: Sagittal slice with hippocampal clusters significant at *p* < .05 (familywise error-corrected, small volume for bilateral hippocampus). **Middle**: Tilted axial slice (aligned with body of hippocampus) with hippocampal clusters significant at *p* < .05 (familywise error-corrected, small volume for bilateral hippocampus). **Right**: Axial slice with inferior frontal gyrus clusters significant at *p* < .05 (familywise error-corrected, small volume for bilateral inferior frontal gyrus). Red-yellow gradient represents *t*-score. Blue indicates regions of interest for small volume corrections. IFG = Inferior frontal gyrus.

**Figure 7:**
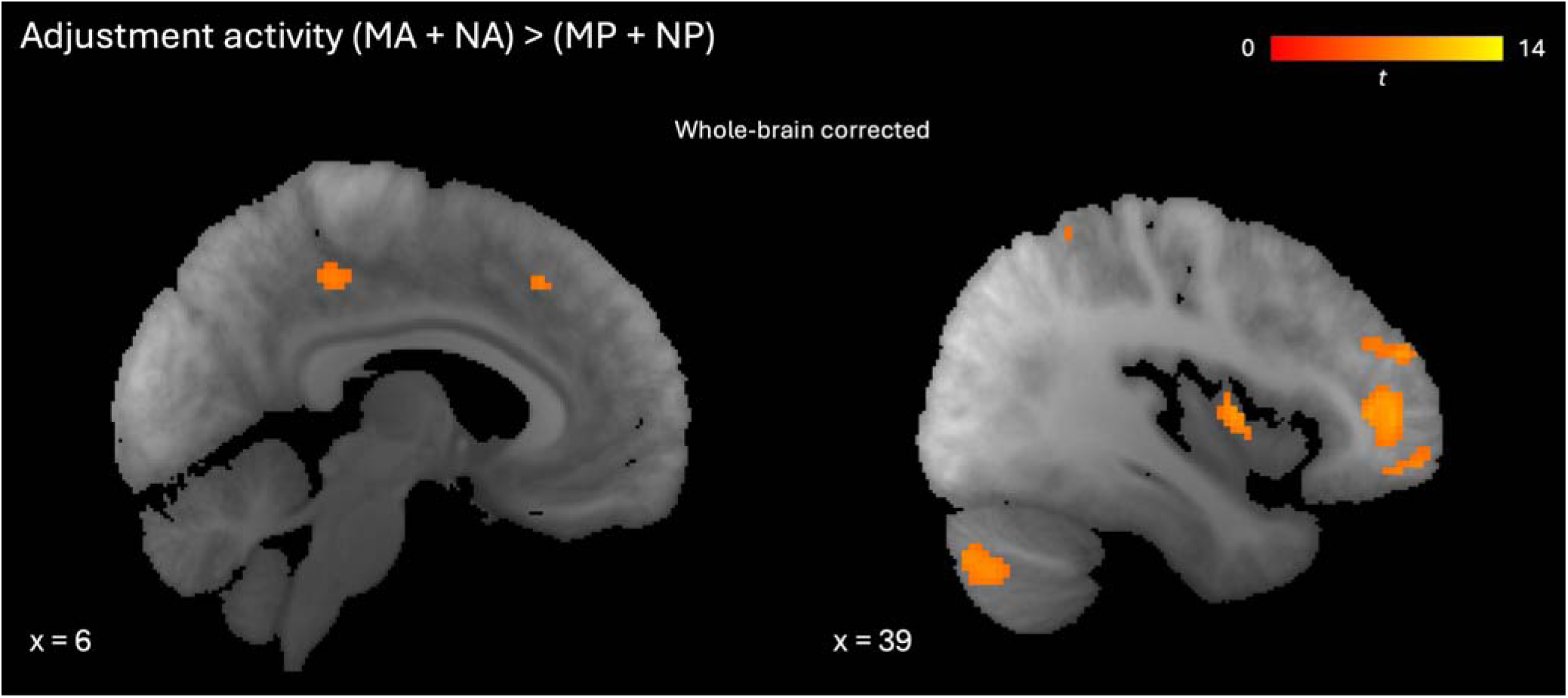
Activity associated with density adjustment in other regions. Clusters showing greater activity for MA and NA conditions than MP and NP conditions significant at *p* <.05 (familywise error-corrected, whole brain). **Left**: Sagittal slice with clusters in posterior cingulate cortex and supplementary motor area. **Right**: Sagittal slice with clusters in frontal pole, insula, and cerebellum. Red-yellow gradient represents *t*-score.

**Table 5.**
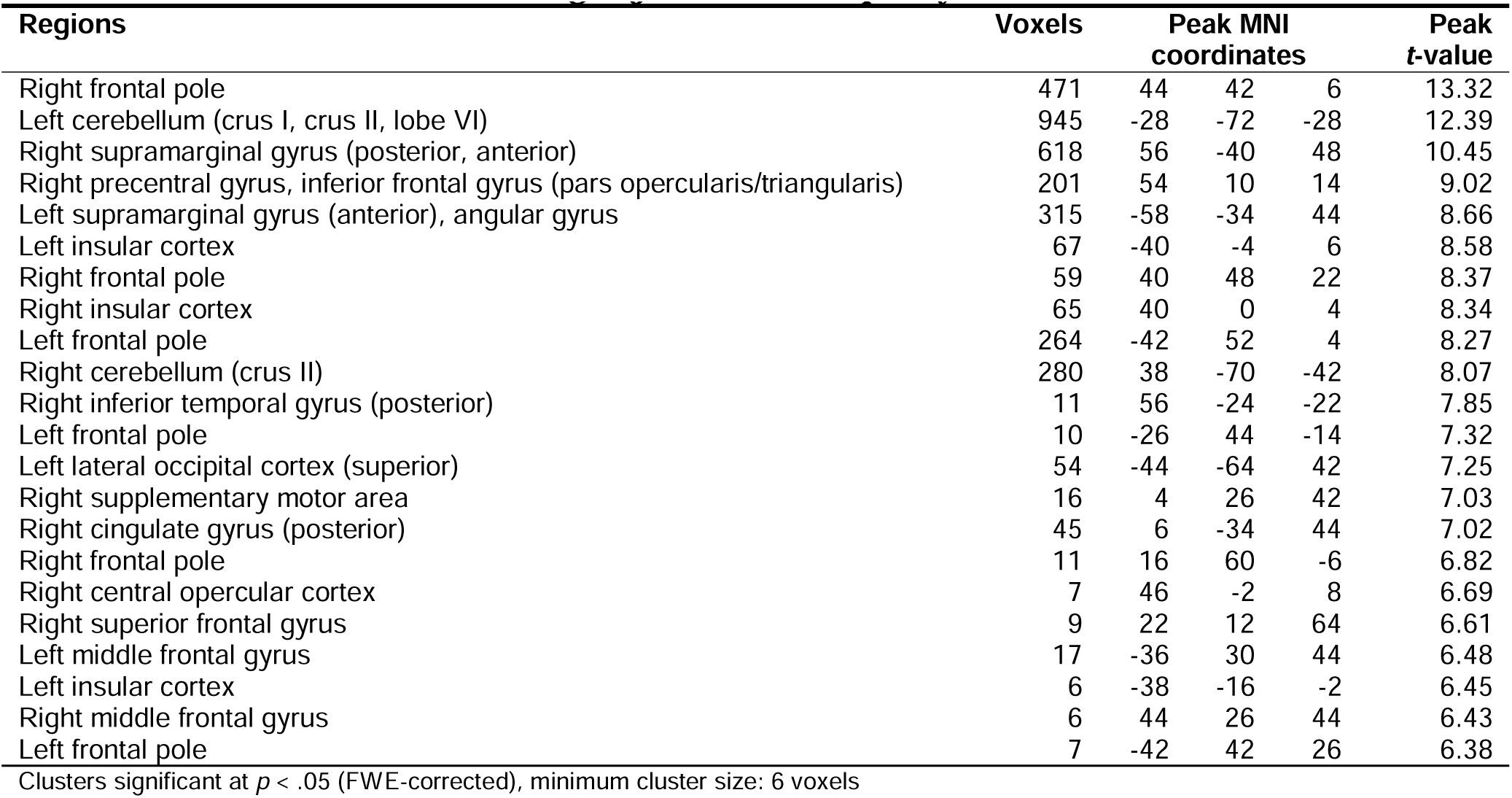
Clusters more active during adjustment than parity conditions.

### Navigation toward a target

We next looked for sites that showed an interaction between memory and adjustment, i.e. for which adjustment-related activity was greatest when there was a target to be navigated toward [(MA-MP)>(NA-NP)]. No regions survived a family-wise error correction at *p* < .05, however at a more liberal threshold (*p* < .001 uncorrected), bilateral orbitofrontal cortex was more active for navigation toward a target than for aimless navigation (Figure 8; Supplemental Table S10). Activity levels for each condition averaged across voxels in each significant cluster are shown in Supplemental Figure 2. This result was qualitatively unaffected by the exclusion of participants reporting spatial associations (Supplemental Table S11). There was no interaction effect in any of our regions of interest.

**Figure 8:**
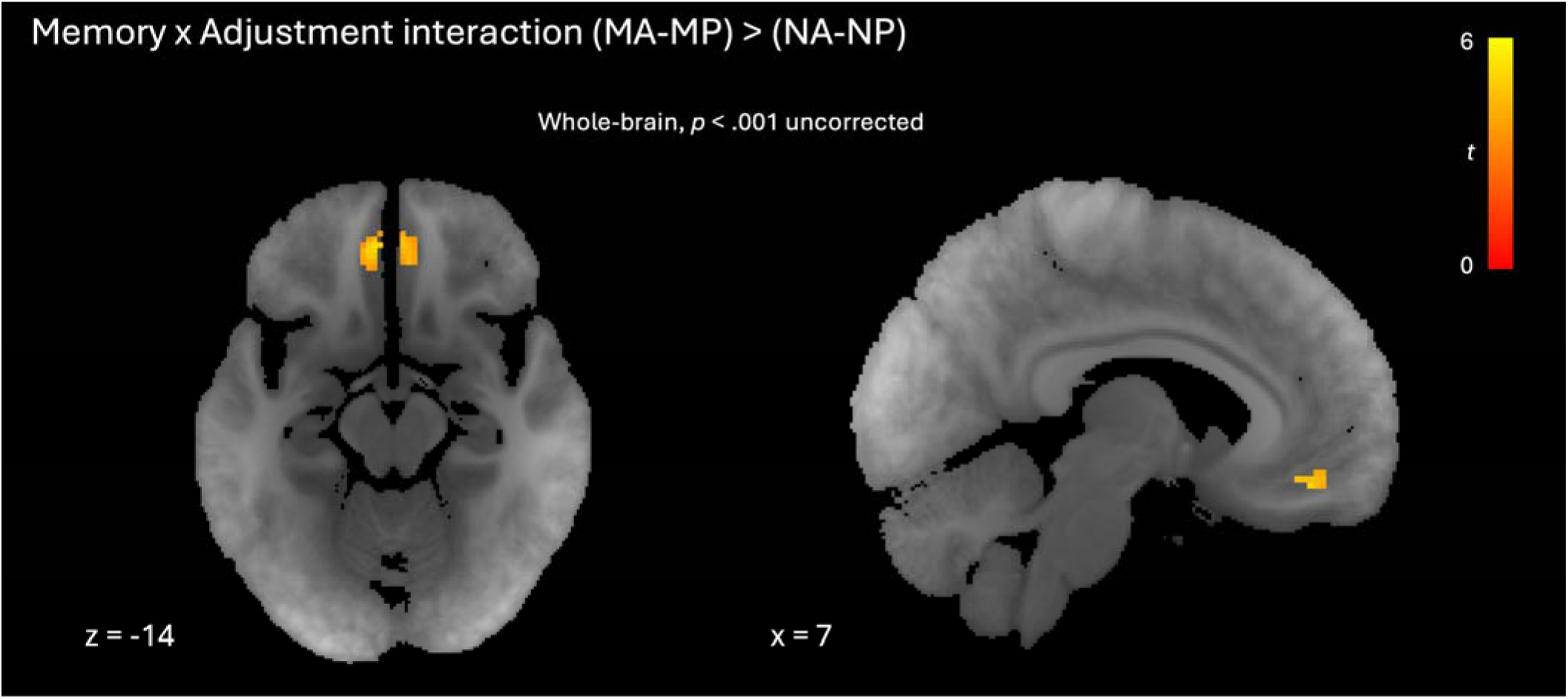
Adjustment activity relating to target presence. Axial and sagittal slices showing interaction between memory and adjustment in orbitofrontal cortex. Clusters significant at *p* < .001 (uncorrected). Red-yellow gradient represents *t*-score.

### Direction and distance from target

No regions showed an encoding of adjustment direction in MA and NA trials. To investigate correlates of distance from target, we had to account for its negative correlation with time in trial. We assessed the unique contribution of distance from target in an analysis in which elapsed time during the adjustment period was entered as an initial regressor, and distance from target as a second orthogonal regressor. This analysis included only the interval in each MA or NA trial during which adjustments actually occurred, along with matching intervals in the corresponding MP and NP trials. No clusters showed effects of distance at the corrected whole brain level, nor did our regions of interest after small volume correction. There were significant clusters at the *p* < .001 uncorrected level in frontal and parietal cortex (Table 6), with the largest cluster in left postcentral gyrus likely reflecting a greater likelihood of button pressing when participants were more distant from the target; this peak was close to that shown in Figure 7B at which adjustment direction was encoded. Effects of elapsed time can be found in Supplemental Tables S12 and S13.

**Table 6.** Clusters with adjustment activity modulated by distance from target (uncorrected).

| Modulation direction | Regions | Voxels | Peak MNI coordinates |  |  | Peak t-value |
| --- | --- | --- | --- | --- | --- | --- |
| Positive | Left postcentral gyrus | 251 | -50 | -22 | 50 | 4.64 |
| Negative | Right supramarginal gyrus | 69 | 60 | -32 | 46 | 4.38 |
|  | Right frontal pole | 45 | 46 | 48 | 8 | 4.29 |
|  | Right postcentral gyrus, precentral gyrus | 48 | 40 | -26 | 60 | 4.18 |

Clusters significant at $p < .001$ (uncorrected), minimum cluster size: 31 voxels

### Navigation performance as a function of encoding and maintenance activity

Navigation to an auditory target requires that the target is first adequately encoded. In a final analysis we looked for brain areas in which encoding and maintenance activity was parametrically associated with performance in the MA trials (i.e. negatively related to the final distance from the target; see Table 7, Figure 9). Two peaks in right auditory cortex (Figure 9 bottom left) were in areas at which the density of the currently heard sound was encoded. Other sites with activity associated with subsequent performance were posterior cingulate gyrus (Figure 9 bottom right) and, after small-volume corrections, hippocampus and inferior frontal gyrus (Figure 9 top, Supplemental Table S14). These effects were specific to the adjustment task: none of these clusters’ encoding/maintenance activity was modulated by performance in the parity task in the MP trials.

**Figure 9:**
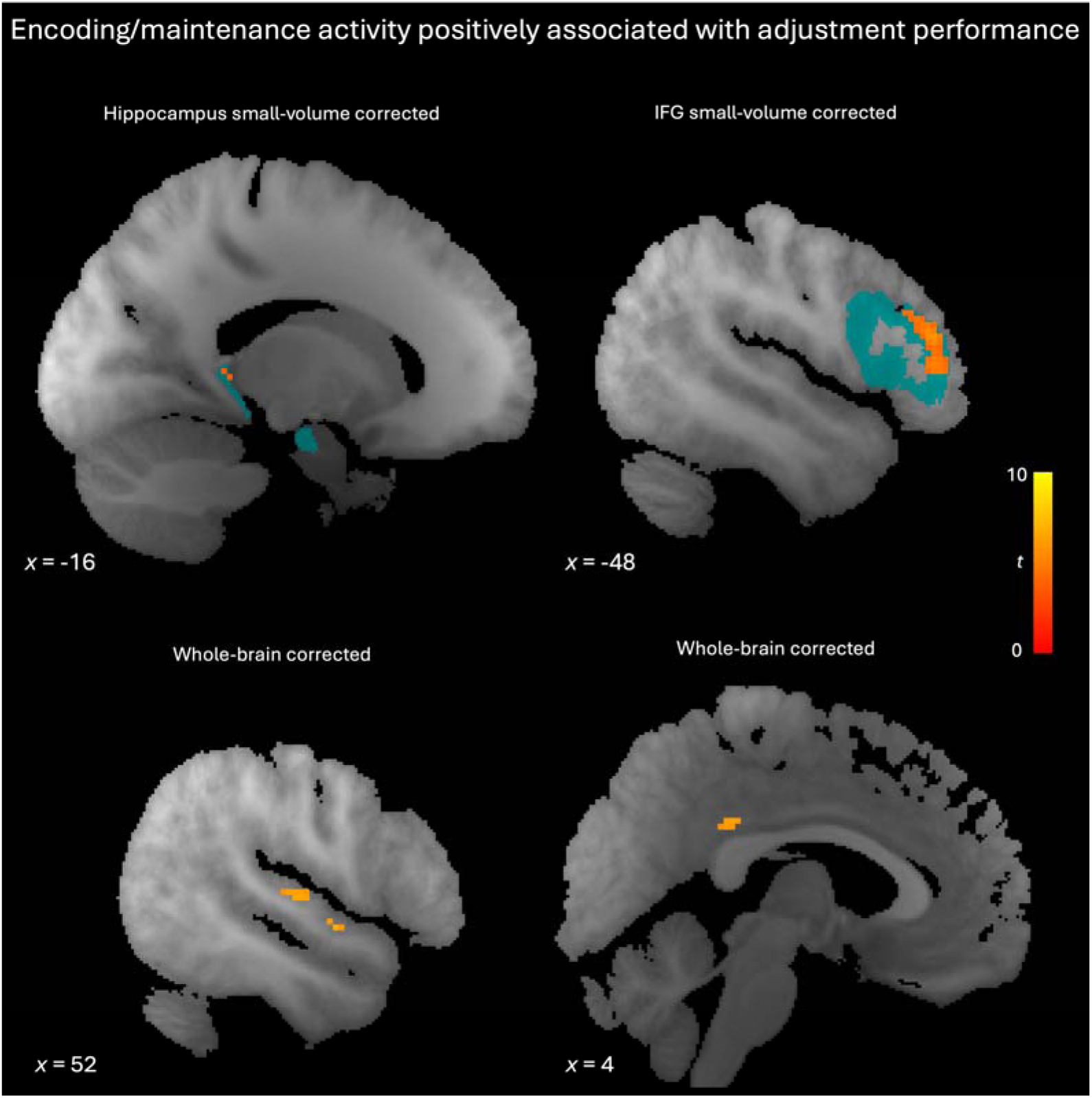
Encoding and maintenance activity positively associated with navigation performance. **Top left:** Sagittal slice with hippocampal cluster significant at *p* < .05 (familywise error-corrected, small volume for bilateral hippocampus). **Top right:** Sagittal slice with inferior frontal gyrus clusters significant at *p* < .05 (familywise error-corrected, small volume for bilateral inferior frontal gyrus). **Bottom left:** Sagittal slice with clusters in auditory cortex (familywise error-corrected, whole brain). **Bottom right:** Sagittal slice with cluster in posterior cingulate cortex (familywise error-corrected, whole brain). Red-yellow gradient represents *t*-score. Blue indicates regions of interest for small volume corrections. IFG = Inferior frontal gyrus.

**Table 7.** Clusters with encoding/maintenance activity positively modulated by adjustment performance.

| Regions | Voxels | Peak MNI coordinates |  |  | Peak t-value |
| --- | --- | --- | --- | --- | --- |
| Right middle frontal gyrus | 17 | -32 | 12 | 60 | 9.26 |
| Right Heschl's gyrus | 43 | 54 | -14 | 4 | 7.32 |
| Right planum temporale | 12 | 62 | -26 | 12 | 7.12 |
| Left cingulate gyrus (posterior) | 16 | -2 | -32 | 30 | 7.03 |
| Right planum polare | 5 | 52 | -2 | -6 | 7.02 |
| Right cingulate gyrus (posterior) | 19 | 2 | -34 | 30 | 6.96 |
| Left occipital pole | 7 | -18 | -100 | 10 | 6.55 |

Clusters significant at $p < .05$ (FWE-corrected), minimum cluster size: 5 voxels

## Discussion

We studied the neural basis for perceiving, storing, and “navigating” toward an auditory target, defined by the density of its acoustic spectrum, a parameter that participants did not consistently map to any physical spatial dimension. It was represented most strongly in non-primary auditory cortex, and adjusting it engaged a broad network including inferior frontal gyrus and hippocampus. Orbitofrontal cortex was differentially activated when adjustments were made to match a target sound, compared to aimless navigation. The density of such target sounds was represented in left inferior temporal gyrus and right anterior hippocampus and the level of activity in auditory cortex, inferior frontal gyrus, hippocampus, and posterior cingulate during target encoding and maintenance was positively associated with navigation performance.

We refer to self-initiated adjustment in this auditory density space as “navigation” but acknowledge that this stretches the standard definition of the term. Our study was partly motivated by suggestions that hippocampus and neighbouring entorhinal cortex, known to be involved in mapping physical space, can also represent task-relevant sensory spaces (Killian et al., 2012; Aronov et al., 2017; Bao et al., 2019). The adjustment-related activity we observed in hippocampus and elsewhere was not driven by “low-level” stimulus features (which were matched across conditions), nor by motor demands or direct spatial associations (which control analyses accounted for). An anterior hippocampal cluster dominated by activity after adjustments finished may reflect greater mind-wandering toward the end of these trials (Zeidman et al., 2016) Posterior hippocampus showed more robust activation; it is involved in retrieving sounds from working memory (Kumar et al., 2016) and in spatial navigation (Kühn and Gallinat, 2014) - for example this region shows experience-dependent structural changes in taxi drivers (Maguire et al., 2000). Previous work also demonstrated structural changes in hippocampus (although more anteriorly) associated with a form of navigation through an auditory space in highly trained piano tuners (Teki et al., 2012). Our study supports hippocampal activation during acoustic navigation in novice participants.

Although the above suggests parallels in the neural basis for spatial and auditory navigation, hippocampal activity patterns did not distinguish sparser from denser sounds as they were heard. Aronov et al. (2017) found individual hippocampal neurons that tiled frequency space when relevant to a rat engaged in a related adjustment task. We may not have had a strong enough hippocampal signal for any instantaneous spectral density coding to be identified with the low temporal resolution of fMRI; other techniques such as single neuron recordings from patients may prove fruitful. An alternative explanation is that hippocampus only represents dimensions readily mapped to physical space, such as frequency. Rusconi et al. (2006) showed that mapping sounds that were higher or lower in frequency or pitch to keys positioned higher or lower on a computer keyboard, respectively, led to quicker responses than under the reverse mapping - even when the task was unrelated to frequency. Under this explanation, spatially tuned machinery in rat hippocampus would have been readily co-opted by space-associated sounds in Aronov et al. (2017). We found no such consistent “dimensional overlap” for our stimulus, although a subset of participants reported spatial associations. Another possibility is that neurons in Aronov et al. (2017) were tracking progress toward a goal. Frequencies were always presented in the same order, although their rate of change varied. The observed tuning could in fact reflect firing of hippocampal “time cells” that adapt their temporal tuning scale in accordance with task demands (Macdonald et al., 2011, Shimbo et al., 2021). This argument is reinforced by recent findings that apparent frequency-specific firing in hippocampus or entorhinal cortex is absent in trials in which animals fail to stop at the target sound (Nguyen et al., 2024; Zutshi et al., 2025). We deliberately chose trial-specific target and start densities, such that participants’ trajectories varied over trials, with a view to identifying tuning specific to the acoustic feature. Density tuning was most robust in planum polare but also present in planum temporale and Heschl’s gyrus. It was driven by stronger responses to sparser sounds (chords with individually identifiable frequency components) than denser (more noise-like) ones. This difference might be due in part to greater adaptation within frequency channels for denser tone stacks. Planum polare is also known to reflect higher-order acoustic features, such as pitch change (Warren and Griffiths, 2003) and spectrotemporal correlation (Overath et al., 2008), and to respond to music over scrambled musical sounds (Angulo-Perkins and Concha, 2019).

Whereas previous work found changes in activity relating to working memory for sound in auditory cortex (e.g. Gaab et al., 2003; Linke et al., 2015; Kumar et al., 2016; Czoschke et al., 2021), our contrast comparing memory to non-memory conditions did not. Our silent maintenance period was much shorter (1-3 s) than in other fMRI studies and not designed to be distinguished from the remainder of the trial. Instead, our analysis encompassed the whole trial, which was dominated by periods of auditory stimulation. Any memory-related activity in auditory cortex may have been swamped by responses to the input, present in both memory and no-memory conditions. We did however find memory-related activity in anterior insula, cerebellum and frontal sites including inferior frontal gyrus, consistent with previous work (Zatorre et al., 1994; Kurth et al., 2010; Kumar et al., 2016; King et al., 2019; Brissenden et al., 2021; Llorens et al., 2023).

The role of hippocampus during working memory maintenance has been hard to pin down (Axmacher et al., 2007; Jenseon and Squire, 2011; Stretton et al., 2012). Recent studies have described both elevated firing during working memory maintenance in hippocampal cells selective for particular concepts (Daume et al., 2024) and reduced broadband high frequency power at hippocampal sites at which spectrotemporal ripples held in working memory can nonetheless be decoded (Uluç et al., 2023). It has been argued that not only persistent firing (Fuster and Alexander, 1971) but also “activity-silent” mechanisms involving changes in synaptic weights (Mongillo et al., 2008; Miller et al., 2018) can support working memory maintenance. Holding a target sound in mind may therefore not require increases in the ensemble activity observed indirectly in the BOLD signal. We were however able to distinguish between sparser and denser target sounds in right anterior hippocampus, as well as left inferior temporal gyrus. We also found a link between the level of posterior hippocampal activity and subsequent success in the auditory navigation task. This contrasts with the finding of Kumar et al. (2016) where participants with better tone working memory showed a reduced hippocampal signal during maintenance (although activity was elevated above baseline at the group level). It is possible that our behavioural effect was dominated by activity during target presentation, rather than in the subsequent short silent maintenance period.

This link between auditory navigation success and encoding/maintenance activity, together with the navigation-related activity and multivoxel representations of the target sound, add to growing evidence for a role of the hippocampus in auditory cognition (Billig et al., 2022), disruption of which may mediate links between hearing loss and dementia (Griffiths et al., 2020). The hippocampus is a highly connected structure and in our task likely formed part of a functional network that incorporated non-primary auditory cortex, in which spectral density was encoded. The posterior cingulate was also more active during encoding and maintenance for trials in which participants were subsequently more successful in reaching the target. Changes in posterior cingulate structure and function have been associated with impaired working memory performance in healthy ageing, schizophrenia, and attention deficit hyperactivity disorder (Leech and Sharp, 2014). In our task this region was also more active during adjustment than parity-judgment trials. A role for this region in navigation has been proposed based on (1) indirect fMRI evidence for posterior cingulate grid cell populations (Doeller et al., 2010; Constantinescu et al., 2016) and (2) an association between cingulo-entorhinal connectivity and navigation success (Coughlan et al., 2020). Regions of medial frontal cortex appear to be essential when rats (Basu et al., 2021) or humans (Ciaramelli, 2008; Spiers, 2008) maintain a goal in physical space. We found clusters in bilateral orbitofrontal cortex that may serve the same purpose during auditory navigation, showing a greater memory effect during the adjustment than the non-adjustment task. Effective connectivity has been demonstrated in humans between auditory cortex and hippocampus on the one hand and orbitofrontal cortex on the other (Rolls et al., 2023). It is another site at which grid-like navigation signals have been detected in the BOLD signal (Constantinescu et al., 2016).

Contrary to one of our hypotheses, the presence of an auditory target did not modulate the amount of adjustment-related activity in hippocampus. This differs from findings from work comparing navigation toward a spatial goal with aimless movement or foraging (Cornwell et al., 2008; Gulli et al., 2019). It is possible that participants in our study set themselves an internal goal in trials in which they were asked to make random adjustments. Directly comparing spatial and auditory navigation correlates in the same participants would also be illuminating.

Distinguishing activity relating to adjustment and working memory processes required a complex design. Rather than comparing conditions involving these processes to a silent baseline, we used a parity-judgment task as a control that minimized hippocampal activity while matching motor activity and sensory input. It required participants to redirect attention within a trial between tone stacks and digits. This had no significant impact on working memory performance (participants stored the target density similarly well in MA and MP trials) but the “adjustment” effects reported should be interpreted with the nature of the baseline task in mind. We also note that some regions, including parts of hippocampus, showed reduced activity in all conditions compared to the inter-trial rest periods. Despite this, the factorial design allowed us to identify condition-specific correlates of auditory adjustment and working memory that existed alongside any general reduction of task-unrelated activity.

### Conclusion

We developed an acoustic stimulus defined by its spectral density, a feature represented in non-primary auditory cortex and not consistently associated with any spatial dimension. Navigation through this density space engaged brain regions including inferior frontal gyrus and hippocampus, with orbitofrontal cortex activated more when navigation was toward a target. Encoding and maintenance in areas including hippocampus, inferior frontal gyrus, auditory cortex and cingulate cortex was positively associated with subsequent successful navigation, and target sounds were encoded in left inferior temporal gyrus and right anterior hippocampus. Overall, we find that self-initiated travel in humans along a pure auditory dimension engages a brain system overlapping that previously found to support spatial navigation.

## Data availability statement

The data that support the findings of this study are openly available in BIDS format at OpenNeuro at https://openneuro.org/datasets/ds006211.

## Conflict of interest disclosure

The authors declare no competing financial interests.

## Acknowledgments

This work was supported by the Wellcome Trust, United Kingdom (WT106964MA), and Medical Research Council, United Kingdom (MR/T032553/1). The Wellcome Centre for Human Neuroimaging was supported by core funding from the Wellcome Trust (203147/Z/16/Z). We thank Oliver Josephs, Nadège Corbin, and Martina Callaghan for assistance with MRI sequence design, preprocessing, and denoising; Eleanor Maguire and Karl Friston for advice on experimental design; and the radiographers at the The Wellcome Centre for Human Neuroimaging (now UCL Department of Imaging Neuroscience).

## Notes

https://openneuro.org/datasets/ds006211

### Competing Interest Statement

The authors have declared no competing interest.

### Summary of Updates

Replaced all within-participant multivariate analyses with across-participant multivariate analyses. Main results results were unchanged, but a representation of the target density was also found in right anterior hippocampus and left inferior temporal gyrus. Other minor changes to figures and discussion.

## References

Ahveninen J, Uluç I, Raij T, Nummenmaa A, Mamashli F (2023) Spectrotemporal content of human auditory working memory represented in functional connectivity patterns. Commun Biol 6:294. 10.1038/s42003-023-04675-8

Angulo-Perkins A, Concha L (2019) Discerning the functional networks behind processing of music and speech through human vocalizations. PLoS One 14:e0222796. 10.1371/journal.pone.0222796

Aronov D, Nevers R, Tank, DW (2017) Mapping of a non-spatial dimension by the hippocampal–entorhinal circuit. Nat 543:719–722. 10.1038/nature21692

Axmacher N, Mormann F, Fernandez G, Cohen MX, Elger CE, Fell J (2007) Sustained neural activity patterns during working memory in the human medial temporal lobe. J Neurosci 27:7807–7816. 10.1523/JNEUROSCI.0962-07.2007

Bao X, Gjorgieva E, Shanahan LK, Howard, JD, Kahnt T, Gottfried JA (2019) Grid-like neural representations support olfactory navigation of a two-dimensional odor space. Neuron 102:1066–1075. 10.1016/j.neuron.2019.03.034

Basu R, Gebauer R, Herfurth T, Kolb S, Golipour Z, Tchumatchenko T, Ito HT (2021) The orbitofrontal cortex maps future navigational goals. Nat 599:449–452. 10.1038/s41586-021-04042-9

Bicanski A, Burgess N (2020) Neuronal vector coding in spatial cognition. Nat Rev Neurosci 21:453–470. 10.1038/s41583-020-0336-9

Billig AJ, Lad M, Sedley W, Griffiths, TD (2022) The hearing hippocampus. Prog Neurobiol 218:102326. 10.1016/j.pneurobio.2022.102326

Brissenden JA, Tobyne SM, Halko MA, Somers DC (2021) Stimulus-specific visual working memory representations in human cerebellar lobule VIIb/VIIIa. J Neurosci 41: 1033–1045. 10.1523/JNEUROSCI.1253-20.2020

Ciaramelli, E (2008) The role of ventromedial prefrontal cortex in navigation: A case of impaired wayfinding and rehabilitation. Neuropsychologia 46:2099–2105. 10.1016/j.neuropsychologia.2007.11.029

Constantinescu, AO, O’Reilly JX, Behrens TEJ (2016) Organizing conceptual knowledge in humans with a gridlike code. Science 352:1464–1468. 10.1126/science.aaf0941

Corbin N, Todd N, Friston KJ, Callaghan MF (2018) Accurate modeling of temporal correlations in rapidly sampled fMRI time series. Hum Brain Mapp 39:3884–3997. 10.1002/hbm.24218

Cornwell BR, Johnson LL, Holroyd T, Carver FW, Grillon C (2008) Human hippocampal and parahippocampal theta during goal-directed spatial navigation predicts performance on a virtual Morris water maze. J Neurosci 28:5983–5990. 10.1523/JNEUROSCI.5001-07.2008

Coughlan G, Zhukovsky P, Puthusseryppady V, Gillings R, Minihane A-M, Cameron D, Hornberger M (2020). Functional connectivity between the entorhinal and posterior cingulate cortices underpins navigation discrepancies in at-risk Alzheimer’s disease. Neurobiol Aging 90:110–118. 10.1016/j.neurobiolaging.2020.02.007

Czoschke S, Fischer C, Bahador T, Bledowski C, Kaiser J (2021) Decoding concurrent representations of pitch and location in auditory working memory. J Neurosci 41:4658–4666. 10.1523/JNEUROSCI.2999-20.2021

Daume J, Kamiński J, Salimpour Y, Gómez Palacio Schjetnan A, Anderson WS, Valiante TA, Mamelak AN, Rutishauser U (2024) Persistent activity during working memory maintenance predicts long-term memory formation in the human hippocampus. Neuron 112:1–12. 10.1016/j.neuron.2024.09.013

Deutsch P, Czoschke S, Fischer C, Kaiser J, Bledowski C (2023) Decoding of working memory contents in auditory cortex is not distractor-resistant. J Neurosci 43:3284–3293. 10.1523/JNEUROSCI.1890-22.2023

Doeller CF, Barry C, Burgess N (2010) Evidence for grid cells in a human memory network. Nat 463: 657–661. 10.1038/nature08704

Ekstrom AD, Kahana MJ, Caplan JB, Fields TA, Isham EA, Newman EL, Fried I (2003) Cellular networks underlying human spatial navigation. Nat 425:184–188. 10.1038/nature01964

Fuster JM, Alexander GE (1971) Neuron activity relating to short-term memory. Science 173:652–654. 10.1126/science.173.3997.652

Gaab N, Gaser C, Zaehle T, Jancke L, Schlaug G (2003) Functional anatomy of pitch memory - an fMRI study with sparse temporal sampling. Neuroimage 19:1417–1426. 10.1016/S1053-8119(03)00224-6

Griffiths TD, Lad M, Kumar S, Holmes E, McMurray B, Maguire EA, Billig AJ, Sedley W (2020) How can hearing loss cause dementia? Neuron, 108:401–412. 10.1016/j.neuron.2020.08.003

Gulli RA, Duong LR, Corrigan BW, Doucet G, Williams S, Fusi S, Martinez-Trujillo JC (2020) Context-dependent representations of objects and space in the primate hippocampus during virtual navigation. Nat Neurosci 23:103–112. 10.1038/s41593-019-0548-3

Hafting T, Fyhn M, Molden S, Moser M-B, Moser EI (2005) Microstructure of a spatial map in the entorhinal cortex. Nat 436:801–806. 10.1038/nature03721

Hebart MN, Görgen K, Haynes J-D (2015) The Decoding Toolbox (TDT): a versatile software package for multivariate analyses of functional imaging data. Front Neuroinform 8:88. 10.3389/fninf.2014.00088

Jeneson A, Squire, LR (2011) Working memory, long-term memory, and medial temporal lobe function. Learn Mem 19:15–25. 10.1101/lm.024018.111

Killian NJ, Jutras MJ, Buffalo EA (2012) A map of visual space in the primate entorhinal cortex. Nat 491:761–764. 10.1038/nature11587

King M, Hernandez-Castillo CR, Poldrack RA, Ivry RB, Diedrichsen J (2019) Functional boundaries in the human cerebellum revealed by a multi-domain task battery. Nat Neurosci 22:1371–1378. 10.1038/s41593-019-0436-x

Kumar S*, Gander PE*, Berger JI*, Billig AJ, Nourski KV, Oya H, Kawasaki H, Howard MA, Griffiths, TD (2021) Oscillatory correlates of auditory working memory examined with human electrocorticography. Neuropsychologia 150:107691. 10.1016/j.neuropsychologia.2020.107691

Kumar S, Joseph S, Gander PE, Barascud N, Halpern AR, Griffiths TD (2016) A brain system for auditory working memory. J Neurosci 36:4492–4505. 10.1016/j.neuropsychologia.2020.107691

Kurth F, Zilles K, Fox PT, Laird AR, Eickhoff SB (2010) A link between the systems: Functional differentiation and integration within the human insula revealed by meta-analysis. Brain Struct Funct 214:519–534. 10.1007/s00429-010-0255-z

Kühn S, Gallinat J (2014) Segregating cognitive functions within hippocampal formation: A quantitative meta analysis on spatial navigation and episodic memory. Hum Brain Mapp 35:1129–1142. 10.1002/hbm.22239

Leech R, Sharp DJ (2014) The role of the posterior cingulate cortex in cognition and disease. Brain 137: 12–32. 10.1093/brain/awt162

Linke AC, Cusack R (2015) Flexible information coding in human auditory cortex during perception, imagery, and STM of complex sounds. J Cog Neurosci 27:1322–1333. 10.1162/jocn_a_00780

Llorens A, Bellier L, Blenkmann, AO, Ivanovic J, Larsson PG, Lin JJ, Endestad T, Solbakk A-K, Knight RT (2023) Decision and response monitoring during working memory are sequentially represented in the human insula. iScience 26:107653. 10.1016/j.isci.2023.107653

MacDonald CJ, Lepage KQ, Eden UT, Eichenbaum, H (2011) Hippocampal “time cells” bridge the gap in memory for discontiguous events. Neuron 71: 737–749. 10.1016/j.neuron.2011.07.012

Maguire EA, Gadian DG, Johnsrude IS, Good CD, Ashburner J, Frackowiak RSJ, Frith CD (2000) Navigation-related structural change in the hippocampi of taxi drivers. Proc Natl Acad Sci U S A 97: 4398–4403. 10.1073/pnas.070039597

Miller EK, Lundqvist M, Bastos AM (2018). Working memory 2.0. Neuron 100: 463–475. 10.1016/j.neuron.2018.09.023

Mongillo G, Barak O, Tsodyks M (2008) Synaptic theory of working memory. Science 319: 1543–1546. 10.1126/science.1150769

Nguyen D, Wang G, Wafa T, Fitzgerald T, Gu Y (2024) The medial entorhinal cortex encodes multisensory spatial information. Cell Rep 43: 114813. 10.1016/j.celrep.2024.114813

Nozari N, Martin, RC (2024) Is working memory domain-general or domain-specific? Trends Cogn Sci, 28:1023–1036. 10.1016/j.tics.2024.06.006

O’Keefe J, Dostrovsky J (1971) The hippocampus as a spatial map. Preliminary evidence from unit activity in the freely-moving rat. Brain Res 34:171–175. 10.1016/0006-8993(71)90358-1

Overath T, Kumar S, Von Kriegstein K, Griffiths TD (2008) Encoding of spectral correlation over time in auditory cortex. J Neurosci 28:13268–13273. 10.1523/JNEUROSCI.4596-08.2008

Pomper U, Curetti LZ, Chait M (2023) Neural dynamics underlying successful auditory short-term memory performance. Eur J Neurosci 58:3859–3878. 10.1111/ejn/16140

Rolls ET, Deco G, Huang C-C, Feng J (2023) The human orbitofrontal cortex, vmPFC, and anterior cingulate cortex effective connectome: Emotion, memory, and action. Cereb Cortex 33:330–356. 10.1093/cercor/bhac070

Rusconi E, Kwan B, Giordano B, Umilta C, Butterworth B (2006) Spatial representation of pitch height: The SMARC effect. Cognition 99:113–129. 10.1016/j.cognition.2005.01.004

Shimbo A, Izawa E-I, Fujisawa S (2021) Scalable representation of time in the hippocampus. Sci Adv 7:eabd7013. 10.1126/sciadv.abd7013

Spiers HJ (2008) Keeping the goal in mind: Prefrontal contributions to spatial navigation. Neuropsychologia 46:2106–2108. 10.1016/j.neuropsychologia.2008.01.028

Stark CEL, Squire LR (2001) When zero is not zero: The problem of ambiguous baseline conditions in fMRI. Proc Natl Acad Sci U S A 98:12760–12766. 10.1073/pnas.221462998

Stretton J, Winston G, Sidhu M, Centeno M, Vollmar C, Bonelli S, Symms M, Koepp M, Duncan JS, Thompson PJ (2012) Neural correlates of working memory in Temporal Lobe Epilepsy—An fMRI study. Neuroimage 60:1696–1703. 10.1016/j.neuroimage.2012.01.126

Tavares RM, Mendelsohn A, Grossman Y, Williams CH, Shapiro M, Trope Y, Schiller D (2015) A map for social navigation in the human brain. Neuron 87: 231–243. 10.1016/j.neuron.2015.06.011

Teki S*, Kumar S*, von Kriegstein K, Stewart L, Lyness CR, Moore BCJ, Capleton B, Griffiths TD (2012) Navigating the auditory scene: An expert role for the hippocampus. J Neurosci 32: 12251–12257. 10.1523/JNEUROSCI.0082-12.2012

Uluç I, Peled N, Paulk AC, Bush A, Gumenyuk V, Kotlarz P, Lankinen K, Mamashli F, Matsuda N, Richardson MR, Stufflebeam SM, Cash SS, Ahveninen J (2023) Decoding auditory working memory content from intracranial high frequency activity in humans. bioRxiv 552073 10.1101/2023.08.04.552073

Uluç I, Schmidt TT, Wu Y-H, Blankenburg F (2018) Content-specific codes of parametric auditory working memory in humans. Neuroimage 183:254–262. 10.1016/j.neuroimage.2018.08.024

Wang Q, Cagna B, Chaminade T, Takerkart S (2020) Inter-subject pattern analysis: A straightforward and powerful scheme for group-level MVPA. Neuroimage 204:116205. 10.1016/j.neuroimage.2019.116205

Warren JD, Griffiths TD (2003) Distinct mechanisms for processing spatial sequences and pitch sequences in the human auditory brain. J Neurosci 23:5799–5804. 10.1523/JNEUROSCI.23-13-05799.2003

Whittington JCR, Muller TH, Mark S, Chen G, Barry C, Burgess N, Behrens TEJ (2020) The Tolman-Eichenbaum Machine: unifying space and relational memory through generalization in the hippocampal formation. Cell 183:1249–1263. 10.1016/j.cell.2020.10.024

Yushkevich PA, Pluta JB, Wang H, Xie L, Ding SL, Gertje EC, Mancuso L, Kliot D, Das SR, Wolk DA (2014) Automated volumetry and regional thickness analysis of hippocampal subfields and medial temporal cortical structures in mild cognitive impairment. Hum Brain Mapp 36:258–287. 10.1002/hbm.22627

Zatorre R, Evans A, Meyer E (1994) Neural mechanisms underlying melodic perception and memory for pitch. J Neurosci 14: 1908–1919. 10.1523/JNEUROSCI.14-04-01908.1994

Zeidman P, Maguire E (2016) Anterior hippocampus: the anatomy of perception, imagination and episodic memory. Nat Rev Neurosci 17:173–182. 10.1038/nrn.2015.24

Zutshi I, Apostolelli A, Yang W, Zheng Z, Dohi T, Balzani E, Williams AH, Savin C, Buzsáki G (2025) Hippocampal neuronal activity is aligned with action plans. Nature 649:153–161 10.1038/s41586-024-08397-7

