## Supporting Information for "Brain bases for navigating acoustic features"

Supplemental Table S1	Clusters with activity negatively modulated by density
Supplemental Table S2	Clusters with multivoxel representation of density after	small volume correction
Supplemental Table S3	Clusters more active during memory than non-memory	conditions after small volume correction
Supplemental Table S4	Clusters with multivoxel representation of target density
Supplemental Table S5	Clusters with multivoxel representation of target density after	small volume correction
Supplemental Table S6	Clusters more active during adjustment than parity conditions	after small volume correction
Supplemental Table S7	Clusters more active during adjustment than parity conditions	after small volume correction, excluding participants reporting	spatial associations
Supplemental Table S8	Clusters more active during adjustment than parity conditions	after small volume correction, excluding participants reporting	spatial associations
Supplemental Table S9	Clusters more active during adjustment than parity conditions	after small volume correction, active adjustment period only
Supplemental Table S10	Clusters showing memory by adjustment interaction effect (uncorrected)
Supplemental Table S11	Clusters showing memory by adjustment interaction effect (uncorrected),	excluding participants reporting spatial associations
Supplemental Table S12	Clusters with activity modulated by time in active adjustment period
Supplemental Table S13	Clusters with activity modulated by time in active adjustment period	after small volume correction
Supplemental Table S14	Clusters with encoding/maintenance activity positively modulated by	adjustment performance after small volume correction

Supplemental Figure S1	Trajectories through density space
Supplemental Figure S2	Activity by condition for significant clusters

Supplemental Videos	Example trials from each condition demonstrating what participants	would have seen and heard, based in part on their specific responses.	An inset window is included specifying key	trial parameters as they	change (this information is for the reader and was not shown to	participants).
Filenames:
MA_Example_Trial_With_Sound.mpeg
MA_Example_Trial_With_Sound.mpeg
MA_Example_Trial_With_Sound.mpeg
MA_Example_Trial_With_Sound.mpeg

**Supplemental Table S1. Clusters with activity negatively modulated by density**

| **Regions** | **Voxels** | **Peak MNI coordinates** | | | **Peak**  ***t*-value** |
| --- | --- | --- | --- | --- | --- |
| Right planum polare, Heschl's gyrus | 551 | 56 | 4 | -2 | 16.83 |
| Left Heschl's gyrus | 404 | -46 | -26 | 10 | 10.73 |
| Right insular cortex | 6 | 38 | 2 | -4 | 6.85 |
| Clusters significant at *p* < .05 (FWE-corrected), minimum cluster size: 6 voxels |  |  |  |  |  |

**Supplemental Table S2. Clusters with multivoxel representation of current density after small volume correction**

| **Regions** | **Voxels** | **Peak MNI coordinates** | | | **Peak**  ***t*-value** |
| --- | --- | --- | --- | --- | --- |
| Right planum polare, Heschl's gyrus, superior temporal gyrus (anterior), temporal pole, planum temporale, central opercular cortex | 358 | 56 | 6 | 0 | 8.89 |
| Left planum polare, Heschl's gyrus, superior temporal gyrus (anterior), planum temporale | 301 | -44 | -20 | -4 | 8.00 |
| Right central opercular cortex | 1 | 52 | 4 | 0 | 7.92 |
| Right superior temporal gyrus (posterior) | 10 | 46 | -22 | -2 | 7.06 |
| Left superior temporal gyrus (posterior), middle temporal gyrus (posterior) | 51 | -68 | -24 | 2 | 6.85 |
| Right superior temporal gyrus (posterior) | 55 | 68 | -24 | 6 | 6.83 |
| Left planum temporale | 11 | -58 | -32 | 12 | 6.71 |
| Right planum temporale | 7 | 52 | -30 | 14 | 5.73 |
| Left planum polare | 4 | -48 | -2 | -10 | 5.42 |
| Right superior temporal gyrus (posterior) | 1 | 50 | -10 | -10 | 5.40 |
| Left Heschl's gyrus | 1 | -44 | -26 | 6 | 5.14 |
| Right superior temporal gyrus (anterior) | 1 | 54 | -8 | -8 | 4.95 |
| Right superior temporal gyrus (anterior) | 1 | 54 | -4 | -10 | 4.95 |
| Right inferior frontal gyrus (pars opercularis) | 8 | 58 | 10 | 2 | 5.80 |
| Right inferior frontal gyrus (pars opercularis) | 1 | 52 | 8 | 6 | 4.70 |
| Clusters significant at *p* < .05 (FWE-corrected for small volumes in regions of interest) |  |  |  |  |  |

**Supplemental Table S3. Clusters more active during memory than non-memory conditions after small volume correction**

| **Regions** | **Voxels** | **Peak MNI coordinates** | | | **Peak**  ***t*-value** |
| --- | --- | --- | --- | --- | --- |
| Left inferior frontal gyrus (pars opercularis) | 264 | -48 | 12 | 24 | 6.16 |
| Right inferior frontal gyrus (pars opercularis) | 67 | 48 | 16 | 20 | 5.31 |
| Left inferior frontal gyrus (pars triangularis) | 18 | -50 | 32 | -4 | 5.23 |
| Right inferior frontal gyrus (pars opercularis) | 5 | 56 | 16 | 30 | 4.99 |
| Right inferior frontal gyrus (pars triangularis) | 1 | 52 | 32 | 22 | 4.86 |
| Right inferior frontal gyrus (pars triangularis) | 2 | 44 | 30 | 20 | 4.77 |
| Right inferior frontal gyrus (pars opercularis) | 1 | 48 | 14 | 9 | 4.50 |
| Left inferior frontal gyrus (pars triangularis) | 1 | -42 | 28 | 22 | 4.44 |
| Clusters significant at *p* < .05 (FWE-corrected for small volumes in regions of interest) |  |  |  |  |  |

**Supplemental Table S4. Clusters with multivoxel representation of target density**

| **Regions** | **Voxels** | **Peak MNI coordinates** | | | **Peak**  ***t*-value** |
| --- | --- | --- | --- | --- | --- |
| Left inferior temporal gyrus (anterior) | 10 | -48 | 0 | -42 | 8.26 |
| Clusters significant at *p* < .05 (FWE-corrected), minimum cluster size: 2 voxels |  |  |  |  |  |

**Supplemental Table S5. Clusters with multivoxel representation of target density after small volume correction**

| **Regions** | **Voxels** | **Peak MNI coordinates** | | | **Peak**  ***t*-value** |
| --- | --- | --- | --- | --- | --- |
| Right hippocampus | 4 | 18 | -10 | -16 | 4.66 |
| Clusters significant at *p* < .05 (FWE-corrected for small volumes in regions of interest) |  |  |  |  |  |

**Supplemental Table S6. Clusters more active during adjustment than parity conditions after small volume correction**

| **Regions** | **Voxels** | **Peak MNI coordinates** | | | **Peak**  ***t*-value** |
| --- | --- | --- | --- | --- | --- |
| Right parietal opercular cortex | 11 | 60 | -22 | 16 | 5.98 |
| Right planum polare, central opercular cortex | 8 | 56 | 6 | 0 | 5.69 |
| Left planum polare | 1 | -38 | -24 | 0 | 5.53 |
| Right parietal opercular cortex | 2 | 56 | -32 | 26 | 5.31 |
| Right planum polare | 11 | 52 | -4 | 0 | 5.20 |
| Left planum polare | 1 | -40 | -22 | -2 | 4.88 |
| Right planum polare | 1 | 58 | -6 | 4 | 4.65 |
| Right frontal pole, inferior frontal gyrus (pars triangularis) | 66 | 48 | 36 | 8 | 9.44 |
| Right precentral gyrus, inferior frontal gyrus (pars opercularis, pars triangularis), frontal opercular cortex | 374 | 54 | 10 | 14 | 9.02 |
| Left inferior frontal gyrus (pars triangularis), frontal pole | 84 | -50 | 36 | 12 | 6.45 |
| Left precentral gyrus, inferior frontal gyrus (pars opercularis) | 16 | -54 | 8 | 20 | 5.15 |
| Left inferior frontal gyrus (pars opercularis) | 4 | -50 | 10 | -2 | 4.94 |
| Left inferior frontal gyrus (pars triangularis), frontal pole | 10 | -50 | 30 | -6 | 4.90 |
| Left inferior frontal gyrus (pars triangularis) | 3 | -54 | 22 | 2 | 4.59 |
| Right frontal pole | 2 | 54 | 36 | -2 | 4.42 |
| Right hippocampus | 86 | 18 | -8 | -20 | 5.96 |
| Right hippocampus, parahippocampal gyrus (posterior) | 50 | 20 | -30 | -8 | 5.67 |
| Left hippocampus, parahippocampal gyrus (posterior) | 62 | -22 | -34 | -8 | 4.91 |
| Left hippocampus | 6 | -20 | -14 | -18 | 4.23 |
| Left hippocampus | 1 | -30 | -22 | -20 | 4.02 |
| Clusters significant at *p* < .05 (FWE-corrected for small volumes in regions of interest) |  |  |  |  |  |

**Supplemental Table S7. Clusters more active during adjustment than parity conditions after small volume correction, excluding participants reporting spatial associations**

| **Regions** | **Voxels** | **Peak MNI coordinates** | | | **Peak**  ***t*-value** |
| --- | --- | --- | --- | --- | --- |
| Right inferior frontal gyrus (pars opercularis) | 87 | 58 | 12 | 14 | 7.29 |
| Right frontal pole, inferior frontal gyrus (pars triangularis) | 8 | 48 | 36 | 8 | 6.48 |
| Right inferior frontal gyrus (pars opercularis) | 3 | 58 | 12 | 2 | 5.96 |
| Right inferior frontal gyrus (pars triangularis) | 1 | 50 | 18 | -4 | 5.77 |
| Right hippocampus | 6 | 16 | -12 | -18 | 7.71 |
| Right parahippocampal gyrus (posterior), hippocampus | 7 | 18 | -30 | -10 | 5.81 |
| Clusters significant at *p* < .05 (FWE-corrected for small volumes in regions of interest) |  |  |  |  |  |

**Supplemental Table S8. Clusters more active during adjustment than parity conditions after small volume correction, excluding participants with mismatched button presses**

| **Regions** | **Voxels** | **Peak MNI coordinates** | | | **Peak**  ***t*-value** |
| --- | --- | --- | --- | --- | --- |
| Right frontal pole, inferior frontal gyrus (pars triangularis) | 29 | 48 | 36 | 8 | 8.82 |
| Right inferior frontal gyrus (pars opercularis), precentral gyrus | 213 | 56 | 10 | 14 | 8.21 |
| Left precentral gyrus, inferior frontal gyrus (pars opercularis) | 7 | -54 | 8 | 22 | 5.32 |
| Left inferior frontal gyrus (pars triangularis) | 4 | -50 | 36 | 12 | 5.28 |
| Left inferior frontal gyrus (pars triangularis), orbitofrontal cortex | 2 | -50 | 30 | -6 | 5.09 |
| Right hippocampus, parahippocampal gyrus (anterior) | 63 | 20 | -14 | -24 | 6.07 |
| Clusters significant at *p* < .05 (FWE-corrected for small volumes in regions of interest) |  |  |  |  |  |

**Supplemental Table S9. Clusters more active during adjustment than parity conditions after small volume correction, actual adjustment period only**

| **Regions** | **Voxels** | **Peak MNI coordinates** | | | **Peak**  ***t*-value** |
| --- | --- | --- | --- | --- | --- |
| Right central opercular cortex, planum polare | 1 | 58 | 4 | 2 | 4.70 |
| Right precentral gyrus, inferior frontal gyrus (pars opercularis) | 204 | 54 | 10 | 14 | 6.20 |
| Left frontal pole, inferior frontal gyrus (pars triangularis) | 23 | -48 | 38 | 14 | 5.25 |
| Right frontal pole, inferior frontal gyrus (pars triangularis) | 6 | 46 | 36 | 10 | 4.58 |
| Left parahippocampal gyrus (posterior) | 10 | -22 | -32 | -20 | 5.96 |
| Right hippocampus, parahippocampal gyrus (posterior) | 1 | 20 | -30 | -8 | 5.67 |
| Right hippocampus, parahippocampal gyrus (posterior) | 1 | 22 | -34 | -8 | 4.91 |
| Left parahippocampal gyrus (anterior) | 1 | -16 | -22 | -20 | 4.02 |
| Clusters significant at *p* < .05 (FWE-corrected for small volumes in regions of interest) |  |  |  |  |  |

**Supplemental Table S10. Clusters showing memory by adjustment interaction effect (uncorrected)**

| **Regions** | **Voxels** | **Peak MNI coordinates** | | | **Peak**  ***t*-value** |
| --- | --- | --- | --- | --- | --- |
| Right orbitofrontal cortex | 29 | 4 | 40 | -16 | 5.31 |
| Left orbitofrontal cortex | 28 | -6 | 44 | -14 | 5.14 |
| Clusters significant at *p* < .001 (uncorrected), minimum cluster size: 26 voxels |  |  |  |  |  |

**Supplemental Table S11. Clusters showing memory by adjustment interaction effect (uncorrected), excluding participants reporting spatial associations**

| **Regions** | **Voxels** | **Peak MNI coordinates** | | | **Peak**  ***t*-value** |
| --- | --- | --- | --- | --- | --- |
| Right orbitofrontal cortex | 7 | 6 | 42 | -16 | 6.05 |
| Left orbitofrontal cortex | 3 | -6 | 44 | -16 | 4.56 |
| Clusters significant at *p* < .001 (uncorrected), orbitofrontal cortex only |  |  |  |  |  |

**Supplemental Table S12. Clusters with activity modulated by time in active adjustment period**

| **Modulation direction** | **Regions** | **Voxels** | **Peak MNI coordinates** | | | **Peak**  ***t*-value** |
| --- | --- | --- | --- | --- | --- | --- |
| Positive | Right precentral gyrus | 239 | 34 | -24 | 64 | 8.77 |
|  | Right precentral gyrus | 21 | 22 | -12 | 72 | 8.73 |
|  | Right superior parietal lobule | 195 | 28 | -44 | 70 | 8.17 |
|  | Right parietal operculum | 10 | 40 | -20 | 22 | 7.47 |
|  | Left cerebellum (lobe V) | 31 | -16 | -50 | -22 | 7.41 |
|  | Left lateral occipital cortex (superior) | 22 | -22 | -86 | 28 | 7.17 |
|  | Right lateral occipital cortex (superior), occipital pole | 88 | 28 | -82 | 26 | 7.05 |
|  | Right occipital pole | 10 | 4 | -92 | -8 | 6.92 |
|  | Right supplementary motor area | 15 | 6 | -12 | 52 | 6.59 |
| Negative | Left postcentral gyrus | 117 | -46 | -22 | 60 | 7.26 |
| Clusters significant at *p* < .05 (uncorrected), minimum cluster size: 7 voxels | |  |  |  |  |  |

**Supplemental Table S13. Clusters with activity modulated by time in active adjustment period after small volume correction**

| **Modulation direction** | **Regions** | **Voxels** | **Peak MNI coordinates** | | | **Peak**  ***t*-value** |
| --- | --- | --- | --- | --- | --- | --- |
| Positive | Right inferior frontal gyrus (pars triangularis) | 28 | 50 | 22 | -6 | 5.03 |
|  | Right hippocampus | 2 | 30 | -22 | -20 | 4.53 |
| Negative | Right superior temporal gyrus (posterior), Heschl's gyrus, planum temporale | 55 | 66 | -12 | 2 | 5.71 |
|  | Left planum temporale | 49 | -56 | -20 | 6 | 5.67 |
|  | Right superior temporal gyrus (posterior) | 23 | 66 | -16 | 0 | 5.66 |
|  | Left Heschl's gyrus | 17 | -34 | -26 | 10 | 5.22 |
|  | Right superior temporal gyrus (posterior) | 3 | 64 | -12 | -4 | 4.87 |
|  | Right superior temporal gyrus (posterior) | 1 | 56 | -14 | -4 | 4.85 |
|  | Right superior temporal gyrus (posterior) | 3 | 58 | -12 | -6 | 4.80 |
|  | Right Heschl's gyrus | 1 | 40 | -26 | 12 | 4.60 |
|  | Right Heschl's gyrus | 1 | 38 | -24 | 10 | 4.58 |
| Clusters significant at *p* < .05 (FWE-corrected for small volumes in regions of interest) | |  |  |  |  |  |

**Supplemental Table S14. Clusters with encoding/maintenance activity positively modulated by adjustment performance after small volume correction**

| **Regions** | **Voxels** | **Peak MNI coordinates** | | | **Peak**  ***t*-value** |
| --- | --- | --- | --- | --- | --- |
| Right Heschl's gyrus, planum temporale, planum polare | 251 | 54 | -14 | 4 | 7.32 |
| Left Heschl's gyrus | 61 | -46 | -24 | 6 | 6.14 |
| Right superior temporal gyrus (anterior) | 5 | 54 | -4 | -10 | 5.07 |
| Left Heschl's gyrus | 1 | -46 | -20 | 2 | 4.64 |
| Left inferior frontal gyrus (pars triangularis) | 138 | -46 | 32 | 16 | 6.37 |
| Right inferior frontal gyrus (pars triangularis) | 37 | 44 | 28 | 16 | 5.49 |
| Left inferior frontal gyrus (pars opercularis) | 5 | -44 | 12 | 28 | 4.63 |
| Left hippocampus | 3 | -14 | -40 | 2 | 4.24 |
| Clusters significant at *p* < .05 (FWE-corrected for small volumes in regions of interest) |  |  |  |  |  |

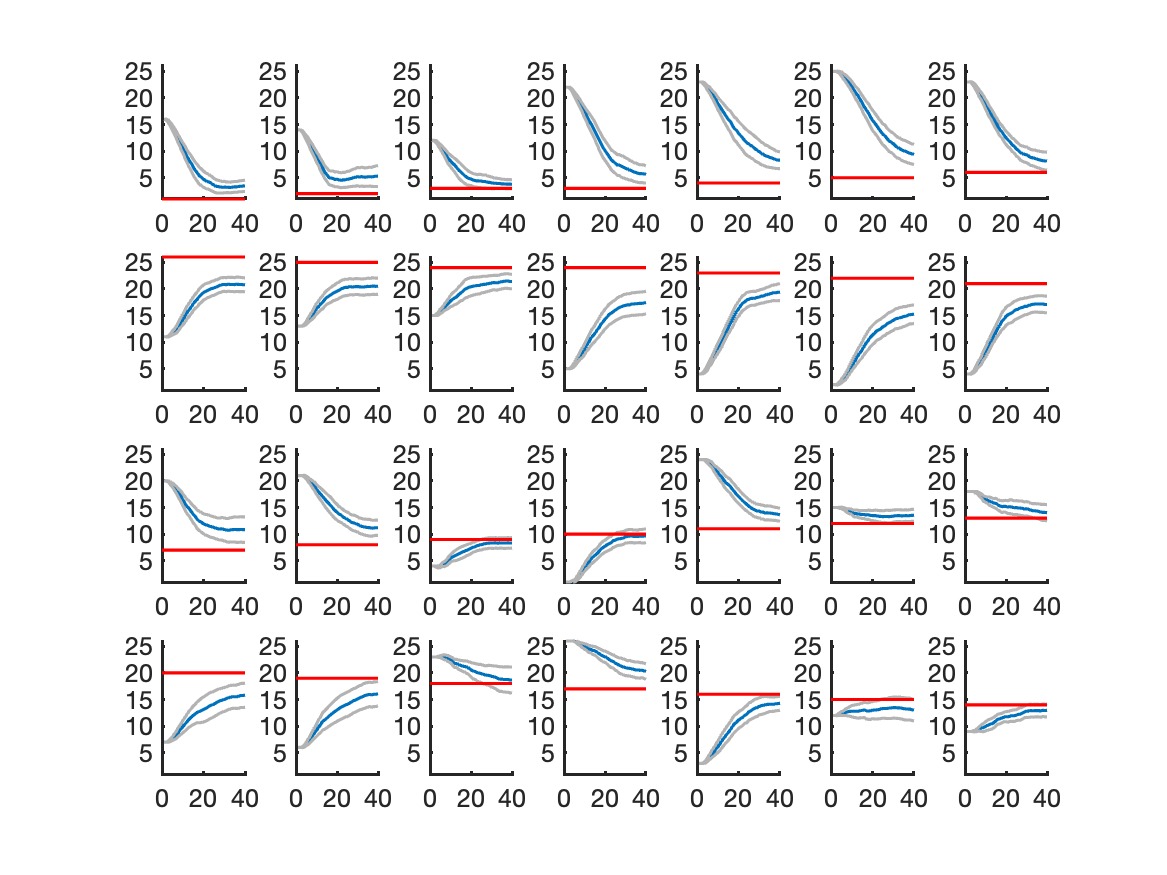

**Supplemental Figure S1: Trajectories through density space.** Each plot shows the mean trajectory (blue) with 95% confidence interval (grey) over participants for a given MA trial, defined by starting density and target density (red line). Stack number (1-40) is shown on the *x*-axis, and density level (1-26, from sparsest to densest) on the *y*-axis. Participants tended to undershoot the target, regardless of whether approaching from sparser or denser sounds.

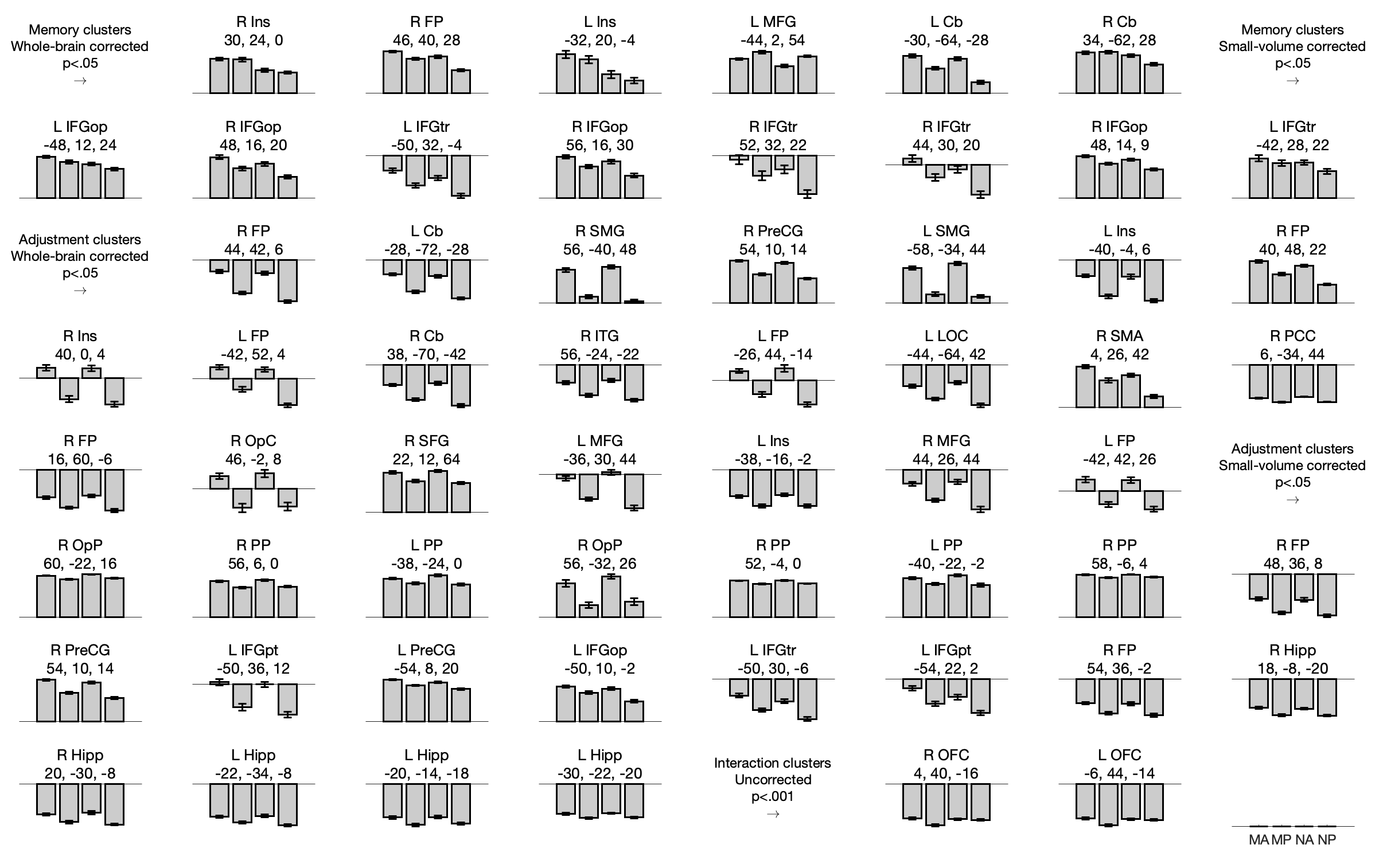

**Supplemental Figure S2: Activity by condition for significant clusters.** Each panel shows activity (in arbitrary units) relative to baseline for each of the four conditions (left to right: MA, MP, NA, NP; see final panel), averaged over all voxels in a given cluster. Error bars show +/- 1 standard error of the mean. All significant clusters for the Memory [(MA+MP)>(NA+NP)], Adjustment [(MA+NA)>(MP+NP)], and Interaction [(MA-MP)>(NA-NP)] effects are shown, as described in the "Memory-related neural activity", "Adjustment-related neural activity", and "Navigation toward a target" sub-sections of Results, respectively. Panels are grouped and ordered as in Tables 4, S3, 5, S4, and S8. Titles give hemisphere, region, and MNI coordinates for the cluster peak. Cb = Cerebellum, FP = Frontal Pole, IFGop = Inferior Frontal Gyrus (pars opercularis), IFGtr = Inferior Frontal Gyrus (pars triangularis), Hipp = Hippocampus, Ins = Insula, ITG = Inferior Temporal Gyrus, L = Left, LOC = Lateral Occipital Cortex, MFG = Middle Frontal Gyrus, OFC = Orbitofrontal Cortex, OpC = Central Operculum, OpP = Parietal Operculum, PCC = Posterior Cingulate Cortex, PP = Planum Polare, PreCG = Precentral Gyrus, R = Right, SFG = Superior Frontal Gyrus, SMA = Supplementary Motor Area, SMG = Supramarginal Gyrus.
